# Welfare of zebra finches used in research

**DOI:** 10.1101/154567

**Authors:** Homare Yamahachi, Anja T. Zai, Ryosuke O. Tachibana, Anna E. Stepien, Diana I. Rodrigues, Sophie Cavé-Lopez, Gagan Narula, Juneseung Lee, Ziqiang Huang, Heiko Hörster, Daniel Düring, Richard H. R. Hahnloser

**Affiliations:** Institute of Neuroinformatics, University of Zurich and ETH Zurich, 8057 Zurich, Switzerland

**Keywords:** zebra finches, songbird, reinforcement, neuronal recording, well-being, stress

## Abstract

Over the past 50 years, songbirds have become a valuable model organism for scientists studying vocal communication from its behavioral, hormonal, neuronal, and genetic perspectives. Many advances in our understanding of vocal learning result from research using the zebra finch, a close-ended vocal learner. We review some of the manipulations used in zebra finch research, such as isolate housing, transient/irreversible impairment of hearing/vocal organs, implantation of small devices for chronic electrophysiology, head fixation for imaging, aversive song conditioning using sound playback, and mounting of miniature backpacks for behavioral monitoring. We highlight the use of these manipulations in scientific research, and estimate their impact on animal welfare, based on the literature and on data from our past and ongoing work. The assessment of harm-benefits tradeoffs is a legal prerequisite for animal research in Switzerland. We conclude that a diverse set of known stressors reliably lead to suppressed singing rate, and that by contraposition, increased singing rate may be a useful indicator of welfare. We hope that our study can contribute to answering some of the most burning questions about zebra finch welfare in research on vocal behaviors.

## 1. Introduction

Basic scientific research, the acquisition of knowledge without immediate technological or potential medical application, has strongly influenced the remarkable progress of human civilization. Basic research provides us with the knowledge we need to understand the world, cure diseases and, in some cases, prevent catastrophes. A prominent indicator of human progress is the massive increase in life expectancy occurring during the last century. Basic scientific research has led to increased hygiene (Stewardson et al., 2011), food availability through plant biotechnology (Kirakosyan et al., 2009), disease prevention (Bennett et al., 2015), and new cancer therapies (Ye, 2016).

Animal experimentation plays a crucial role in producing knowledge and testing theories of living organisms (Thomas, 2006). However, it is the moral duty of scientists to consider the ethical implications of their research and to continuously weigh the benefits of the outcomes of their animal experiments with the potential harm inflicted on the animal. The universally accepted concept of the 3R principles (Replacement, Reduction and Refinement) is used to guide researchers towards efforts that maximize the quality of life of experimental animals, to closely monitor and continually react to signs of distress, and to follow the best statistical practices for reducing the number of animals required for research. A 4th R, for “Responsibility” was recently proposed by the Max Planck Society, which strives for greater research into the “objective determination of sentience, pain, consciousness, and intelligence in the animal kingdom” (Max Planck Society, 2017). In that vein, it is important to assess the burden induced by experimental manipulations of animals, which is the central aim of this paper. In our case, we focus on the zebra finch (*Taeniopygia guttata*), a songbird species that is frequently used in research on sexual behaviors and vocal communication.

### 1.1. Zebra finch as a model organism for studies of complex vocal behaviors

The song-control system of songbirds is one of the most intensely studied neural architectures underlying a complex learned behavior. A wide spectrum of insights have been gained in the past 50 years of birdsong research, triggered mainly by the discoveries that birdsong is culturally learned (Thorpe, 1958) and that the brain has evolved a dedicated network of brain areas for singing (Nottebohm et al., 1976).

Thanks to a growing community of researchers, we have learned much about the links between the brain and song-related behaviors. A key advantage of the natural singing behavior in songbirds is the behavioral richness of song and the sensitive readout it provides about the underlying physiological mechanisms. Such richness is not typically encountered when animals are trained using simple operant conditioning paradigms.

In past years, we have learned that successful song learning critically depends on auditory feedback (Kao et al., 2005; Konishi, 1965; Tumer and Brainard, 2007) and on motor exploration (Olveczky et al., 2005). A growing body of evidence indicates that sleep is of great importance for accurate song copying (Derégnaucourt et al., 2005), and we are beginning to understand the underlying mechanisms (Shank and Margoliash, 2009). Also, technological advances are helping us to uncover group dynamics of vocal interactions, with individual-level resolution (Anisimov et al., 2014; Gill et al., 2015; Stowell et al., 2016). There are remarkable parallels between speech learning and birdsong learning (Bolhuis et al., 2010; Brainard and Doupe, 2002; Lipkind et al., 2013). These parallels extend to molecular and hormonal mechanisms of healthy and diseased vocal behaviors (Brainard and Doupe, 2013; Miller et al., 2008). Moreover, brain transcriptomes have revealed a common genetic basis for vocal learning in humans and in songbirds (Haesler et al., 2004; Pfenning et al., 2014).

In this study, we review not the science of birdsong, but the implied impact on animal welfare. We review behavioral needs of zebra finches and the welfare impact of some of the experimental manipulations used in our research. One of our aims is to evaluate whether singing can be used to read out the degree of stress induced by these manipulations. Wherever possible, we propose refinements for some of the manipulations used in our own research.

### 1.2. Review of stress responses and welfare

Welfare and stress are two related concepts. It has been argued that stress comes from an animal’s evaluation of the outcome of a situation, and that welfare is the state resulting from that evaluation (Veissier and Boissy, 2007). Briefly, if a situation evokes stress, then the welfare is poor.

In practical terms, to determine whether an animal would enter a state of stress is to assess whether it would be deprived of the opportunity to act according to its behavioural needs (Dawkins, 1990). However, this is not an easy assessment to make. Determining what counts as a behavioral “need” for any animal species is a complex problem and is thus highly debated (Dawkins, 1983; Friend, 1989). Some authors argue that the necessary criteria for classifying a behavior as a need are the following: a) that all members of the species must perform said behavior, b) that deprivation of that behavior leads to stress, c) the behavior is generated internally and not by external environmental cues, d) that the lack of opportunity to perform a behavior leads to an accumulation of the intent to perform it (Vestergaard, 1982), and e) that the animal attains reward from performing the behavior and hence is inclined to do so even when faced with obstacles (Boissy et al., 2007; Spruijt et al., 2001).

On a behavioral level, measuring the frequency or intensity of behaviors such as fleeing or flight can indicate the presence of a stressor. However, in some animals including birds, stress responses can present themselves as bouts of freezing. In general, to characterize the behavioral needs of an animal, the stress produced by the absence of those needs, and the behavioral and physiological outcomes of stress are complex issues. In the following sections, we briefly review some of the widely adopted readouts of stress and, if possible, highlight their applicability in zebra finches.

#### Corticosterone

Corticosterone is the major glucocorticoid hormone involved in energy regulation, immune function, and stress response in birds. Stressors (e.g. unpredictable events) on birds trigger the release of corticosterone into the bloodstream. Therefore, a direct measurement of corticosterone can, in principle, indicate the stress level in a bird (Sheriff et al., 2011). Both invasive and non-invasive methods to measure corticosterone are available in birds (Sheriff et al., 2011): blood (Washburn et al., 2002), droppings (Goymann, 2005) and feathers (Romero and Fairhurst, 2016).

Corticosterone levels in blood provides an overview of the stress level at that point in time. The total amount of corticosterone in blood depends on circadian (Rensel et al., 2014; Turriani et al., 2016) and seasonal cycles (Dawson and Howe, 1983), on long-term stressors (e.g. extended periods without food), and on acute stressors (e.g. capturing a bird) (Romero and Romero, 2002). Contrary to corticosterone levels in blood, corticosterone in droppings and feathers is considered a chronic measure of the hormone over long time periods (Goymann, 2005; Romero and Fairhurst, 2016): from several hours for droppings to weeks and months for feathers. For this reason, acute stressors might not be reflected in the measured levels of corticosterone in droppings and feathers.

Brain structure and function is directly influenced by stress hormones (McEwen et al., 2015). Therefore, one needs to be aware that plasma levels of corticosterone might not reflect corticosterone levels in the zebra finch brain (Rensel et al., 2014).

For all these reasons, it is important to choose the corticosterone sampling method that is appropriate for measuring the effect of the stressor one aims to characterize.

Short-term increases of corticosterone may be adaptive in nature (Wingfield and Kitaysky, 2002), for example, by enhancing spatial memory (Saldanha et al., 2000). Also, the permanency of endocrine changes and their role in welfare are are still debated. In the case of songbirds, exposure to corticosterone-increasing stressors during development can have both positive and negative effects on birds when they reach adulthood. Such findings detailed below call for caution when interpreting corticosterone levels as indicators of welfare.

On the one hand, stressors during development can negatively impact fitness (Buchanan et al., 2003, 2004; Henriksen et al., 2011; MacDougall-Shackleton and Spencer, 2012; Spencer et al., 2003; Wada et al., 2015). For example, there is evidence that song complexity is inversely related to developmental stress. Namely, zebra finches raised under nutritional stress sing songs containing fewer syllables per song motif as adults (Spencer et al., 2003). Also, zebra finches raised under nutritional stress develop a smaller song control nucleus HVC than when raised with unrestricted food access (Buchanan et al., 2004).

On the other hand, developmental stress can also have positive adaptive effects on fitness and phenotype (Crino and Breuner, 2015). Fledgling European starlings exposed to higher in ovo corticosterone concentration develop larger wing areas and heavier pectoral muscles resulting in an enhanced flight performance (Chin et al., 2009). Early exposure to stress improves spatial learning and memory in juvenile Japanese quails (Calandreau et al., 2011) and associative learning in adult chickens (Goerlich et al., 2012). Corticosterone exposure during development enhances performance of a novel foraging task (Crino et al., 2014a) and reproductive success (Crino et al., 2014b) in adult zebra finches.

Therefore, the interplay between corticosterone levels during development and long-term consequences of stress needs to be further studied.

#### Skin temperature

A stressor can increase the core body temperature, provoking a decrease of the surface temperature. This phenomenon known as stress-induced hyperthermia could be utilized to measure stress. Indeed, infrared thermography has been used to determine the effect of stressful procedures in birds (Edgar et al., 2013; Herborn et al., 2015; Jerem et al., 2015). Contrary to the stress snapshot of corticosterone, this non-invasive imaging method might provide longitudinal measurements of stress.

#### Body weight

Body weight can be used to assess body condition in birds, especially when multiple measurements are made over the course of a study (Brown, 1996; Dickens et al., 2009b, 2009a).

However, body weight can be affected by multiple factors besides stress. In zebra finches, body weight shows a circadian component that fluctuates as much as 10% during the day (Dall and Witter, 1998). In these birds, body weight is also affected by social hierarchy and its effects on feeding behavior (David et al., 2011).

Although stress causes predominantly weight loss, a few studies in both humans and animal models have found weight gain, supporting the “Comfort food” hypothesis (Harris, 2015).

Measurements of body weight as a readout of stress are not routinely reported in birdsong studies. In Section 3.1, we study the effect of changes in husbandry on body weight.

#### Fault bars in feathers

Stressors (e.g. handling, malnutrition, and disease/parasites) can alter the normal development of feathers, producing the appearance of fault bars, which are deformations perpendicular to the rachis (Jovani and Rohwer, 2017). When we verified the tails and wings of 109 birds in our lab, we found putative fault bars (Fig. 1-1) in only 2 birds. A typical lab environment is an unlikely place to find fault bars.

**Figure 1-1.**
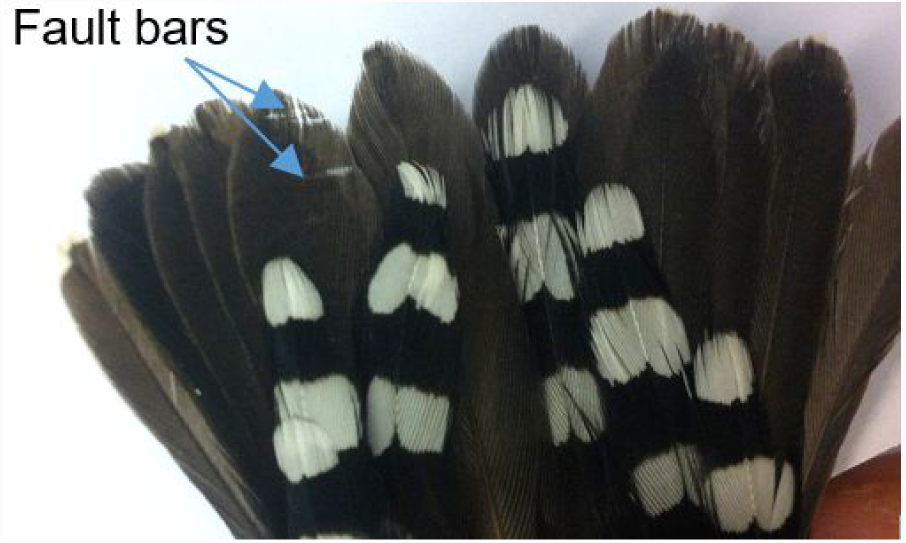
Putative fault bars in the tail feathers of an adult zebra finch housed in a social cage in our lab.

#### Freight molt

Bird species subjected to high degree of predation molt feathers from the rump, back and tail in order to escape (Møller et al., 2006). Therefore, the loss of feathers during a procedure can indicate a stressful event. In our lab, fright molt occurs very rarely during standard husbandry procedures such as checking bird identity, transferring a bird to a new cage, or ringing a bird. In roughly 1 out of 1000 standard procedures a bird may lose a few tail feathers. However, during nonstandard procedures such as bird weighing, fright molt can occur more frequently (in roughly 5/126 ≈ 4% of cases).

### 1.3. General husbandry guidelines

Which environmental factors are important for the welfare of zebra finches?

Housing zebra finches without respecting their natural needs and behaviors can negatively affect their well-being. In the following, we review aspects of husbandry specifically applicable to zebra finches.

#### Housing

Birds should be housed in a physically enriched environment that takes into account their natural environment. Cages should have a variety of features to promote natural behaviors, e.g. space to fly, foraging options, and structural components (Nager and Law, 2010).

In a research facility, cage space is considerably smaller than in zebra finches’ natural habitat. Keeping birds in cages rather than in a large aviary may sometimes be preferable if birds need to be captured and handled on a very regular basis (to eliminate stress for the entire group) (Joint Working Group on Refinement, 2001). Unfortunately, not much is known about the effect of cage size on zebra finches’ behavior (Jacobs et al., 1995). Nonetheless, governmental agencies have set minimum requirements for zebra finches. In Switzerland, the Animal Welfare Ordinance (Tierschutzverordnung, 2008 as of 01.05.2017) states that the minimum space for four estrildid finches (such as zebra finches) must have an area of 0.24 m^2^ with a total volume of 0.12 m^3^. For every additional bird, the floor area must increase by 0.05 m^2^.

#### Feeding and foraging

The main food source of zebra finches in the wild is a variety of seeds. In the cage, besides providing food in containers, a spray of Panicum millet and seeds on the bedding material encourages the display of natural foraging behavior. In addition, cuttlefish bone can be provided to supplement calcium, and egg food to supply proteins in their diet. Besides food, the provision of grit is recommended as a source of minerals, to facilitate the digestion of seeds, and for dust bathing.

#### Water source

Zebra finches inhabit the dry grasslands and desert areas of Australia. These regions have on average 40 days of rain per year and less than 400 mm of rainfall. In captivity, humidity can be kept at 42–55% and ad libitum access to water should be provided in water feeders.

Even though a water source for bathing might be scarce in the wild and zebra finches can survive without water for at least 250 days (Cade et al., 1965), a water bath should be occasionally provided. Indeed, lack of a water bath increases the plasma corticosterone concentration in zebra finches, without affecting body weight (Krause and Ruploh, 2016).

#### Additional enrichment measurements

Perches made out of soft wood or plastic in a range of diameters allow birds to exercise the feet, keep the nails in good condition, and maintain body fitness. To mimic the natural bouncing of branches, it is preferable that perches have a single attachment point. Another type of perch is the swing, on which birds exercise a variety of muscles involved in balancing.

#### Artificial light

Zebra finches can see ultraviolet (UV) light (300–400 nm waveband), so their perception of the environment differs from our own (humans cannot see t below 390 nm) (Cuthill et al., 2000). Zebra finches use their tetrachromatic color space in at least two different situations: mating (Bennett et al., 1996; Hunt et al., 1997) and foraging (Maddocks et al., 2001). In addition, birds are more sensitive to flickering lights than humans, as their flicker fusion threshold is over 100 Hz (Boström et al., 2016; Evans et al., 2012). For these reasons, the light should be flicker free (> 30 kHz), of at least 500 lux, and span the daylight spectrum.

It is important to take light quality into account when designing experiments. Typically, zebra finches should be kept with a constant photoperiod throughout the year (14 hours light to 10 hours dark period, or 12 hours:12 hours). Although seasonal daylight variability has an effect on many birds including reproduction, zebra finches are opportunistic breeders and breed only when conditions are favorable, irrespective of time of year (Small and Moore, 2009).

#### Temperature

Zebra finches adapt to quick increases in temperature, without altering corticosterone levels (Xie et al., 2017). In the wild, temperature can vary drastically during the day and throughout the year, ranging from nearly freezing to over 40 °C. In captivity, a stable temperature between 20 to 26 °C is adequate for both birds and husbandry personnel.

Three particular readouts of stress are reviewed in the next chapter, supplemented by data from our own laboratory: Stress signaled by movement stereotypies, by weight loss, and by reduction or increase in vocalization rate.

## 2. Readout of stress in captive zebra finches

Restrictive conditions in captivity in which animals have little or no control over daily events can lead to stress (Bassett and Buchanan-Smith, 2007). Typically, when animals and humans face negative events, they choose predictability over unpredictability when given a choice (Weinberg and Levine, 1980). Therefore, potentially negative events, such as daily husbandry routines, “should be made temporally predictable” to reduce stress (Bassett and Buchanan-Smith, 2007; Gottlieb et al., 2013).

However, predictability can also lead to boredom (van Rooijen, 1991), so care need to be taken to avoid introducing or modifying husbandry and experimental procedures that could have the effect of increasing stress levels through boredom by not taking into account the species’ natural behaviors (Dawkins, 2004).

In birds, vocalization is the main form of communication with conspecifics (Suzuki, 2016; Todt and Naguib, 2000). In zebra finches, two types of singing behaviours are prevalent depending on the social context: directed and undirected singing (Zann, 1996). When males sing directly to a female (i.e. facing the female) their directed song is faster and has less spectral variability than when they sing alone or not facing any bird (Bischof et al., 1981; Kao et al., 2005; Sossinka and Böhner, 1980). As such, experimental procedures that engage both directed and undirected singing in birds might be considered beneficial.

Cognitive engagement, whose use “engages evolved cognitive skills by providing opportunities to solve problems and control some aspect of the environment” has been proposed as a way to increase the well-being of animals in captivity (Clark, 2011). One approach is “to provide animals with specific, cognitively challenging tasks and measure their performance on the same or different tasks at a later date” (Clark, 2017). In the lab, some of our experiments require birds to solve cognitive tasks, such as to hop from a perch to solve an auditory discrimination task (Narula and Hahnloser, 2015) and to modify the song to solve an interactive playback task (Canopoli et al., 2014).

In the following, we discuss stress readouts that could be routinely applied in laboratory studies of vocal behaviors.

### 2.1. Stereotyped behaviors

Monitoring of movement stereotypies and other abnormal behaviors allows assessing an animal’s welfare (Bateson and Feenders, 2010). Abnormal repetitive behavioral patterns, or stereotypies, such as spot picking (i.e. repeated pecking of a spot on itself or on a cage) and route tracing (i.e. repeated pacing behavior) with no obvious goal or function, can develop in animals in captivity (Bateson and Feenders, 2010; van Hoek and ten Cate, 1998; Keiper, 1969; Mason, 1991). Stereotypies can develop over time and become more rigid and more frequent (Meehan et al., 2004). Stereotypies might be associated with a general disinhibition of behavioral control, leading to perseveration on a range of cognitive tasks (Garner et al., 2003a, 2003b). We separately discuss housing conditions and environmental enrichment for prevention of stereotyped behaviors in Sections 3.1-3.3.

To automatically detect individual variation in behavioral performance is challenging. Several commercial systems for monitoring locomotion in a range of animal species (in particular, rodents) have been developed: e.g. EthoVision (Noldus Information Technology, Netherlands), and IntelliCage (NewBehavior AG, Switzerland). However, movement monitoring is not routinely performed in songbird studies in the laboratory.

#### An automated beak tracking system

To read out movement and possibly detect repetitive behaviors in zebra finches, we have devised an automated movement monitoring system. This system consists of a webcam and a simple image processing algorithm. Zebra finches’ characteristic red (in males) and orange beak (in females) can easily be localized in every image frame. Using continuous beak tracking, non-invasive monitoring of a bird’s activity is possible even in tethered birds (Fig. 2-1). An application of our system to detect the effects of environmental manipulations on movement is described in Section 3.2.

**Figure 2-1.**
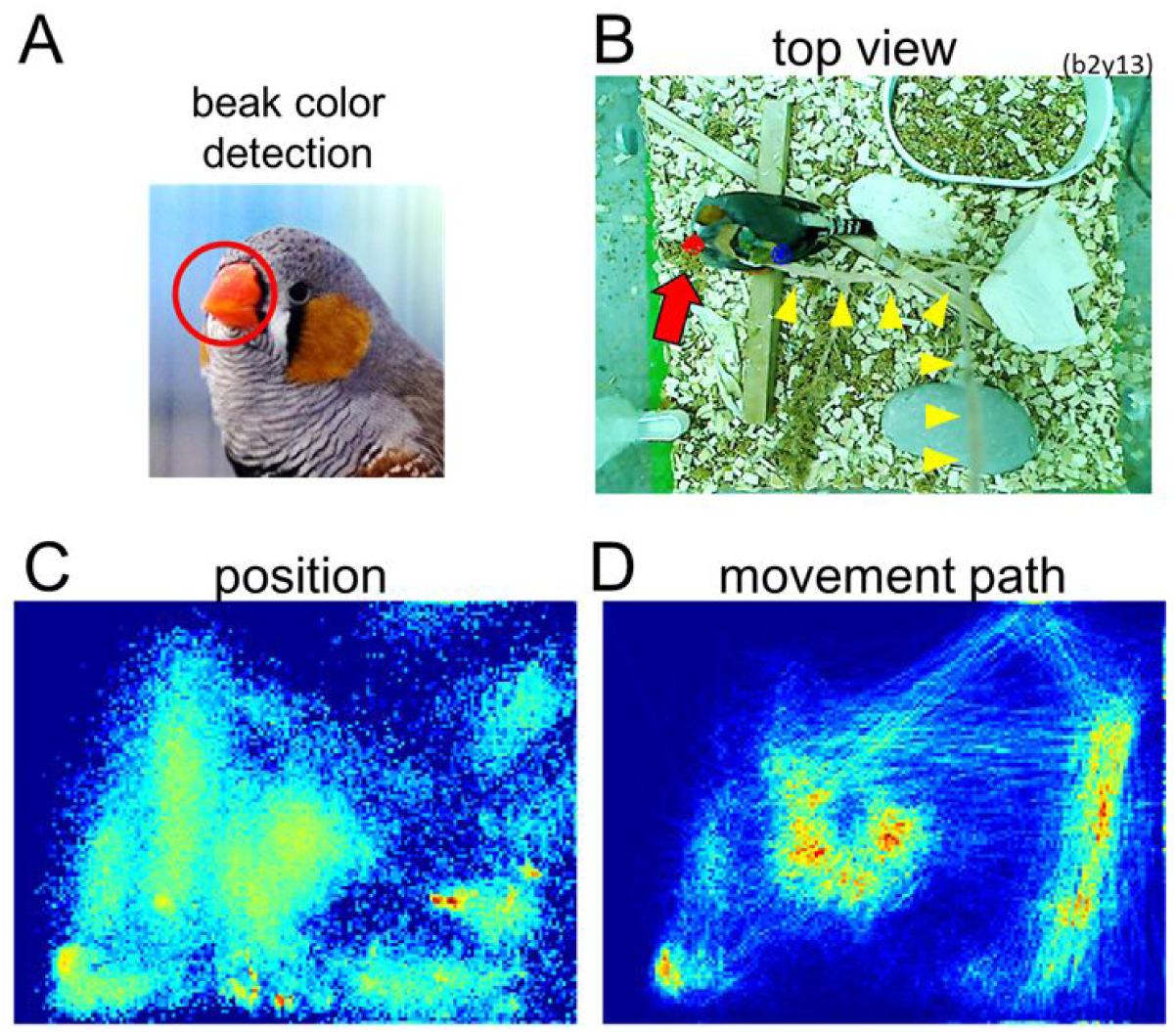
Automatic beak tracking system for movement monitoring. From a continuous video stream (30 frames/s) of the cage interior, we extract the position of the beak (red circle/arrow) based on its hue (A,B). By tracking the beak position we obtain a position map (C), showing the spatial density of the animal, and a movement map (D), showing the movement paths. These maps can be generated from merely 5-minute long recordings. The position map is useful to detect preferred cage locations and longer periods of immobility, and the movement path map can provide information about possible movement stereotypies. Shown are movement data from a 10-hour period obtained in an isolated bird, tethered from the center of the cage. Beak tracking is robust despite the tether (yellow arrowheads) that rapidly moves around in the cage (B).

### 2.2. Calls as a readout of stress

#### Distress calls

Certain calls in zebra finches have been associated with aggressors (Elie and Theunissen, 2016; Zann, 1996). Distress calls are emitted when zebra finches are being attacked or while escaping. A study documented the presence of alarm calls when the experimenter’s hand was ‘in front or in the cage’ (Elie and Theunissen, 2016). In our lab, we have observed distress-like calls when a hand approached a bird or while the cage was being cleaned (Fig. 2-2).

**Figure 2-2.**
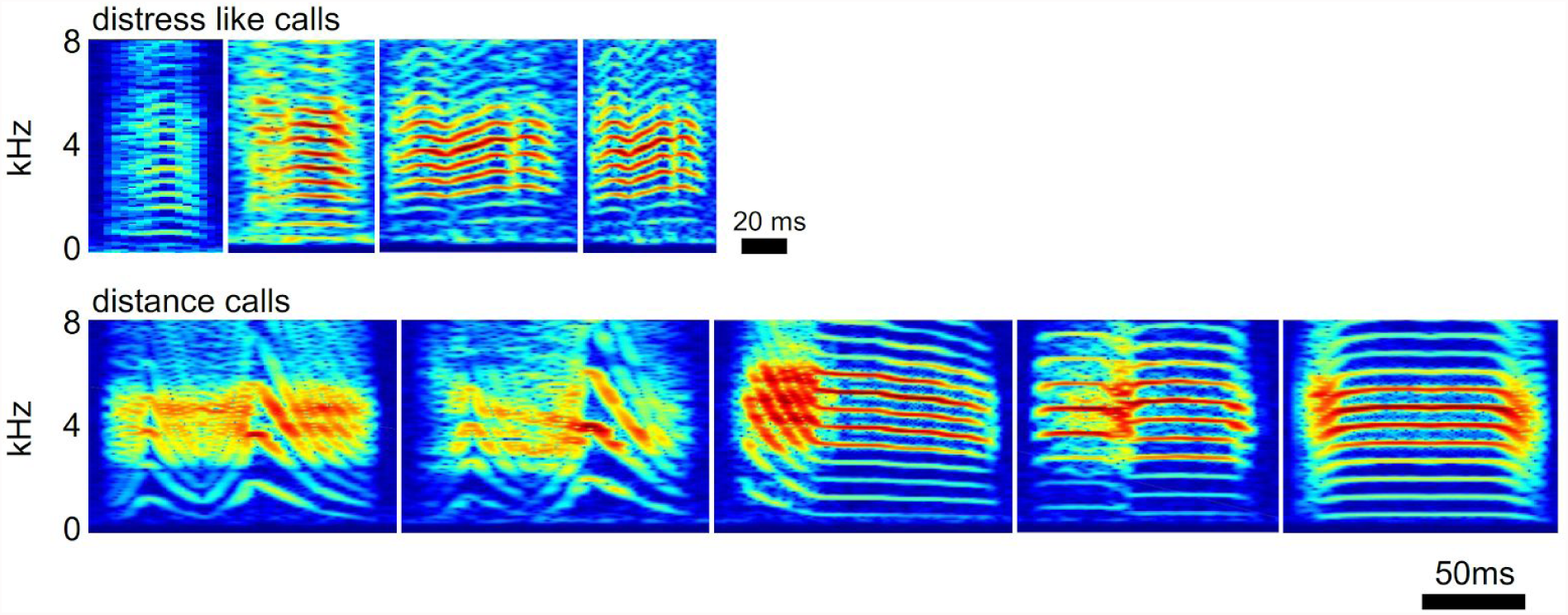
Distress calls and distance calls. Spectrograms of distress-like calls (top panel) emitted when the experimenter’s hand approached the bird (first from left) or while the experimenter cleaned the soundproof box(2nd-4th from left). Spectrograms of learned distance calls (bottom panel) produced by 5 different birds held in isolation.

The conditions under which distress calls are emitted in laboratory environments have not been thoroughly studied. We investigated whether distress calls are emitted in birds that are tethered to a head implant weighing about 1.5 g while they were isolated in a cage of dimensions 24 × 24 × 20 cm. In the span of 10 days each, we found potential distress calls in 2/8 birds only; in one bird we found 13 possible distress calls on one day and 17 on another day. The observed calls were associated with noise in the cage that was caused most likely by the experimenter manipulating or cleaning the cage. In another bird, we found 5 possible distress calls on one day and 5 distress-like calls on another day. All these calls were associated with hand approaches, documented by the experimenter. In other words, on 78/80 days we found no evidence of distress-like calls. Judging by these observations, distress calls are not suitable as a means to read out possible stress induced by our experimental procedures and housing conditions.

#### Distance calls

Zebra finches use distance calls (see Fig 2-2) during long-range communication (Zann, 1996). To our knowledge, the production of distance calls during isolation has not been studied. Birds might use distance calls to try to communicate with other birds. We analyzed data in which birds (*N* = 8) were individually housed initially and then joined together in pairs (*N* = 4).

We found a trend that birds produced more distance calls in isolation than in a social setting (Fig. 2-3). In all bird pairs, the sum of distance calls produced during isolation was higher than the total number of distance calls produced while together. Note that, because of high mutual similarity of distance calls among animals, we did not attempt to assign each distance call to its producer when birds were housed in pairs. Fig. 2-3 also shows that the distance call rates observed during prolonged periods of either isolation or social housing were stable, suggesting that distance call rates do not reflect the time birds have spent in a given social condition.

**Figure 2-3.**
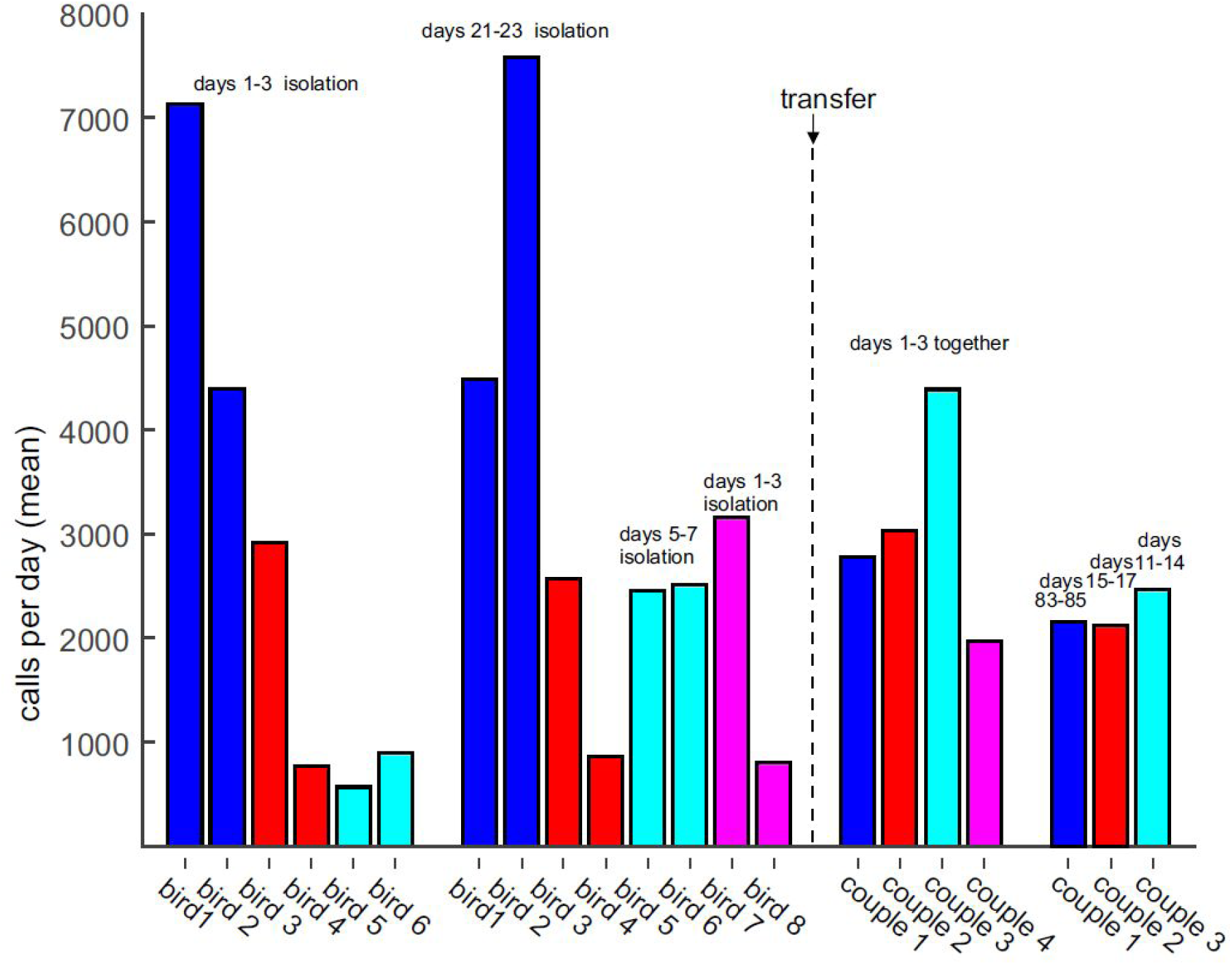
Males produce frequent distance calls in isolation and in a social group. The daily number of distance calls (averaged over 3 consecutive days) is shown in birds that were isolated and then joined in pairs (N = 4 pairs). First, four birds were isolated for 3 weeks (blue and red bars), two birds were isolated for 7 days (cyan bars) and two birds for 3.5 days (purple bars). After joining the birds in pairs, they produced fewer distance calls, shown is the sum of calls produced by each pair. After a long period of social housing (15 days in pair 1, 83 days in pair 2, 11 days pair 3), there was a tendency of small decrease in distance call production (right bars).

There is one caveat to our analysis, which could have affected our interpretation. Namely, birds may have occasionally heard other birds in the lab when the latter were not in soundproof boxes. Such events were not documented, so some of the distance calls recorded could have been produced in exchange with animals not part of the experiment. In other words, a few calls recorded in the first part of the experiment might have been produced not in a state of strict isolation. Despite this caveat, it is obvious that even in a social group, birds produce hundreds of distance calls daily, thus questioning whether these calls can signal a subjective state of isolation.

### 2.3. Singing rate as a readout of welfare

In territorial birds, singing rate can be indicative of diverse properties such as body condition, motivation, parasite load, reproductive success, and territory quality (Collins, 2004; Hall et al., 2009; Searcy and Nowicki, 2005). Song rate is positively correlated with male body condition (Marchetti, 1998; Nyström, 1997), which plays a role in determining the outcome of aggressive interactions between males (Collins, 2004).

Zebra finches are not territorial, and male zebra finches provide paternal care. Female zebra finches in the laboratory prefer males with higher singing rate, and with them they produce heavier offsprings (Collins, 2004; Hall et al., 2009). Thus, in zebra finches too, data indicate that high singing rate might imply good body condition.

The relationship between singing rate and stress has not been studied directly. The prevailing data suggest that singing rate in zebra finches provides a sensitive readout of welfare and that a high singing rate is indicative of good welfare. We arrive at this conclusion by contrapositive logic based on the following two Observations A and B.

Observation A. There is abundant evidence that stress reduces singing rate. 1) Restricted availability of food decrease singing rate (Ritschard and Brumm, 2012). 2) After a surgery (see Fig. 2-4), singing is suppressed between several hours up to a few days (depending on the animal and the surgery). 3) Tethering a bird for the first time, which is a stressor in terms of increased mass and non-zero torque on the head, reduces singing rate for 1–2 days (Fig. 2-5). 4) Isolating a bird in a new environment, a stressor evidenced by a transient increase of corticosterone levels (Banerjee and Adkins-Regan, 2011; Turriani et al., 2016) can decrease singing rate for up to a few days (see Fig. 2-6). 5) Head-fixing a bird tends to suppress song (Picardo et al., 2016). 6) Sick birds do not sing at all. 7) Mounting a backpack on a male zebra finch decreases singing for several days (Anisimov et al., 2014) All these seven observations agree with Selye’s definition of stress (Selye, 1936), according to which diverse aversive manipulations all show the same symptoms (i.e., suppressed singing).

**Figure 2-4.**
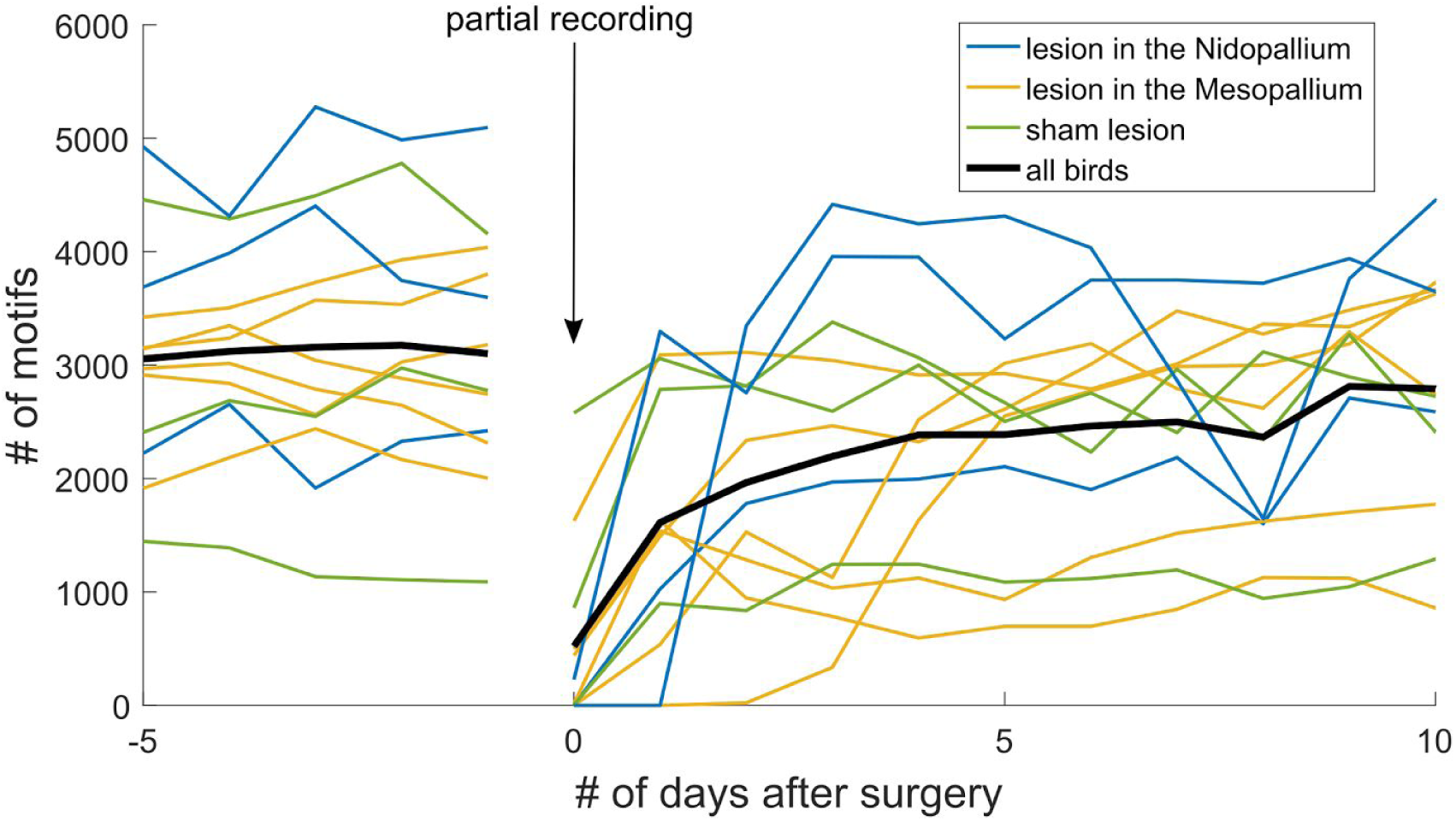
Brain surgeries transiently suppress singing rate. All birds were isolated in a cage (39 × 23 × 39 cm) inside a soundproof box. Shown are the number of song motifs produced from five days before surgery up to 10 days after surgery (N = 12). Bird were given either an injection of saline (green, N = 6) or ibotenic acid (blue and orange, N = 6) delivered into a brain area not controlling song (all surgeries were targeting brain areas outside the song-control system). Because of the surgery, song recording was partial on that day (vertical arrow). Motifs sung before the surgery were not counted, thus the right part of the plot indicates the post-surgery singing rate.

**Figure 2-5.**
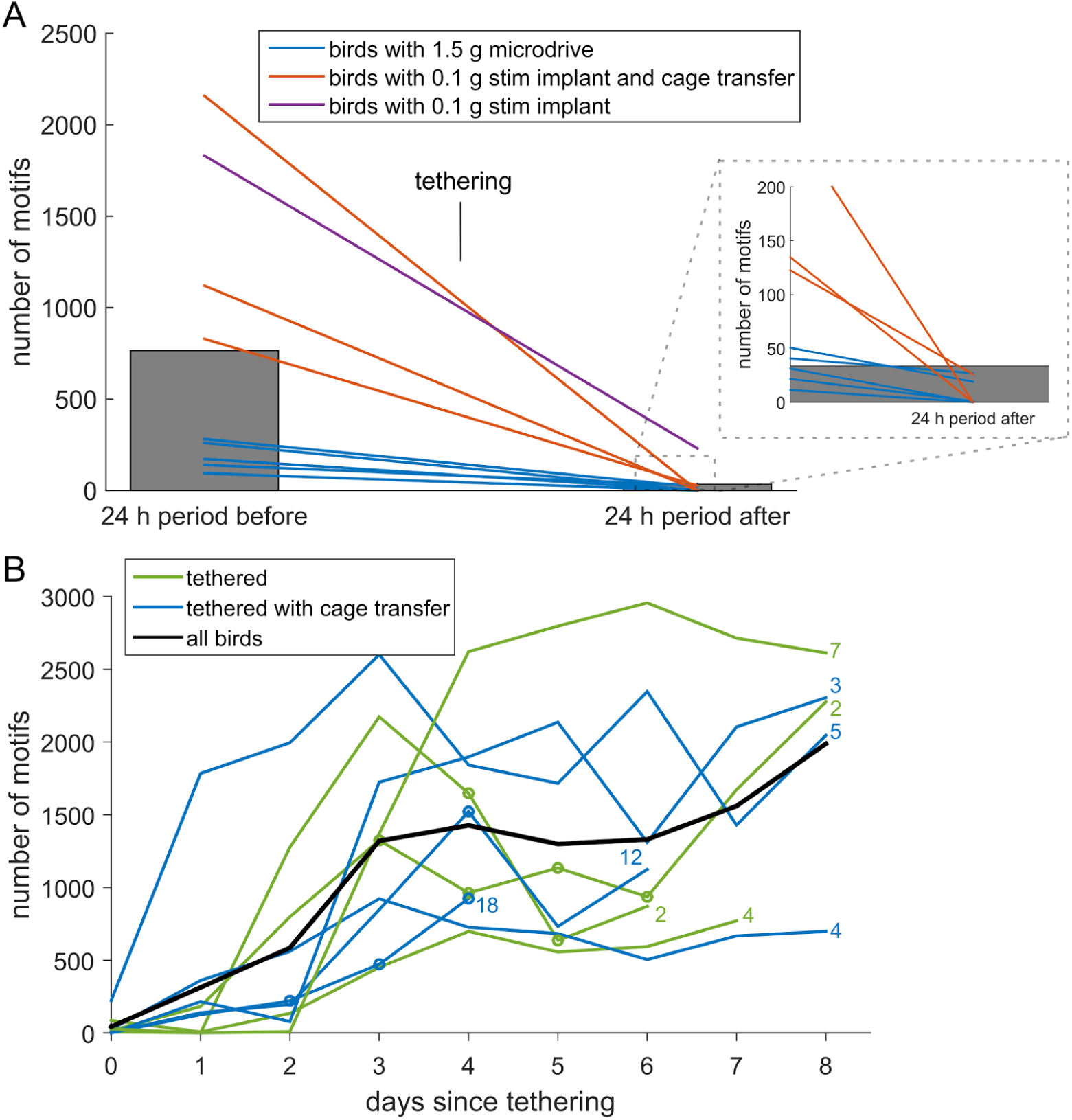
Tethering suppresses singing rate. A) The number of motifs produced time intervals 24 hours before tethering onset (left bar) and 24 hours after tethering onset (continuous tethering, N = 9 birds). All birds were isolated inside a soundproof box. Data in blue (N = 5) were from birds implanted with a 1.5 g microdrive 2–3 days before tethering onset, whereas data in red and violet (N = 4) were from birds implanted with a 0.1 g connector 3–7 days before tethering. In addition, the birds marked in red were transferred to a smaller cage upon tethering. All other birds (blue and violet) were housed in the same cage before and during tethering. Note that the stress reflected by the shown data did not arise from tethering alone, because as indicated in Fig. 2-4, full recovery from surgery can take up to 6 days or more. Hence, the suppressed singing reflects the cumulative burden of surgery and tethering, which is an upper bound to the stress of tethering alone. Note also that it was not possible to extend the analysis past 24 h for birds with microdrive after tethering onset, because birds were subsequently handled. B) Tethering suppresses singing for 1-3 days. Shown is the number of daily song motifs since tethering the animal on day 0. The number of days between the implantation surgery (stimulation device weighing less than 0.5 g) and tethering onset is shown for each animal as a (blue or green) number. Four animals are shown in A, for the others there was no sound recording available before tethering (which is why they are not shown in A).

**Figure 2-6.**
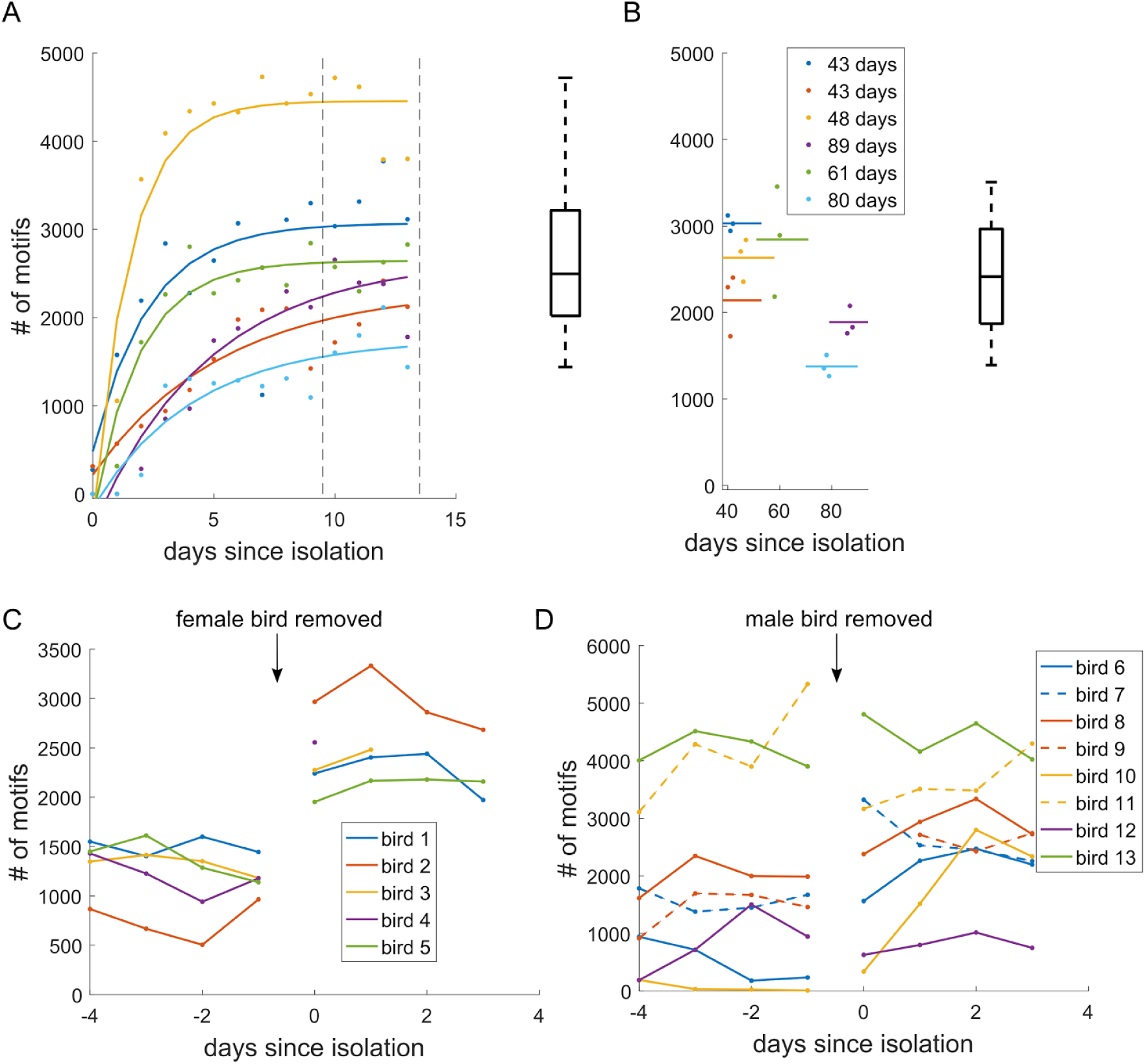
Isolation in a new environment reduces singing rate. A) Number of daily song motifs produced since isolation in a new environment (cage in a soundproof box). The colored dots show data from individual birds (N = 6) and the lines indicate the trend. The boxplots indicate the median, quartiles, and range across birds, computed between days 10 and 14 (dashed vertical lines). B) The daily number of motifs produced after long-term isolation (≥43 days) remains very high (the duration of isolation is indicated in the legend). The lines indicate the daily number of motifs averaged across three consecutive days in the same birds as in A (color-matched lines). The boxplot summarizes the distribution across birds. C, D) Isolating birds in a familiar environment increases singing rate. C) Upon removal of a female social partner (on day 0), the number of motifs per day roughly doubles in the now isolated male (N = 5 birds). D) When two males are separated and isolated in a familiar environment,seven of eight birds strongly increase their daily singing rate. Upon separation, birds 6, 8, 10,12 and 13 stayed in the same soundproof box, the other two birds (7, 9 and 11) were returned to a familiar soundproof box. Data from bird pairs are shown in identical color (dashed line and solid line). Extreme behavior was observed in one couple, in which one bird produced exceptionally many song motifs (dashed yellow), whereas his partner almost never sang (solid yellow). Upon isolation, singing rates became similar in this pair. All these data illustrate that not isolation per se, but isolation in a new environment suppresses singing rate. A-D) Before the isolation event, all birds were first housed for at least 30 days in a same-sex social cage with at least 6 conspecifics.

Observation B. When no obvious stressor is present, zebra finches reliably sing at high rates. In our soundproof boxes, isolated birds sing hundreds of motifs daily. None of our undisturbed bird has ever produced fewer than 400 song motifs per day (we analyzed the songs of 45 birds on more than 1000 days in total; see Fig. 2-7).

**Figure 2-7.**
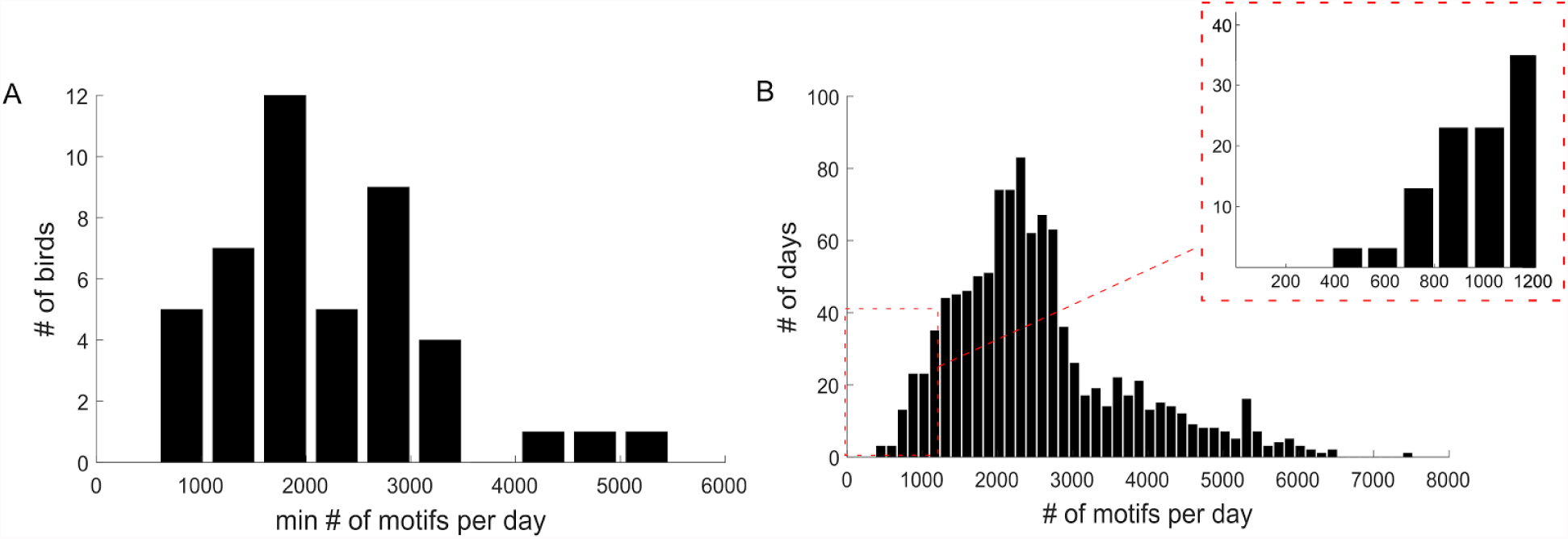
Uninterrupted singing in isolated adults. **A.** Histogram of the minimum number of song motifs produced per day across 45 birds. The minimum in each animal is computed across all days it spent in isolation since familiarization to the new environment. The familiarization period in each animal was at least 5 days long, after which each animal was recorded for at least 12 days. Birds were not subjected to any manipulation except song conditioning (see Section 4.1). **B.** Histogram of the daily number of song motifs produced across all 1066 days analyzed (same 45 adults as in A). No bird ever produced fewer than 400 song motifs per day. A, B)

Contrapositive logic dictates that if A) all clearly stressful manipulations lead to a strong reduction in singing rate, and B) in the absence of a stressor there are no spontaneous events of reduced singing rate, then frequent singing implies absence of stress. In other words, we suggest that frequent singing is a reliable indicator of good welfare. Support for this conclusion is also provided by anecdotal evidence that hand-raised birds tend to sing in the hand of the person who raised them, but birds raised with little human contact do not sing when held.

Our data suggest the presence of a stressor in isolated male zebra finches when they sing less than a certain number of song motifs per day. Ideally, this number should be based on the singing rate the animal displayed before the surgery/procedure, to account for inter-individual variability. Nevertheless, given our observed absolute minimum of 400 song motifs per day in 45 birds, a reasonable number that minimizes false positive detection could be 400 daily song motifs. If we pursue this idea and take a threshold of maximally 400 daily song motifs from Fig. 2-7 as an indicator of stress, we arrive at the conclusion that surgery causes stress in 7/12 animals (Fig. 2-4), isolation in a new environment causes stress in 3/6 animals (Fig. 2-6A), isolation in a familiar environment causes stress in 0/12 animals (Fig. 2-6B,C), and the combination of surgery and tethering causes cumulative stress in 14/14 animals (Fig. 2.5A,B).

Note that none of the examined stressors chronically suppressed singing for more than 4 days (at most 2 days in case of isolation to new environment and at most 4 days after a surgery). Could it perhaps be that singing is such a strong behavioral need that when suppressed for more than 4 days, it becomes a life-threatening condition, similar to lack of bowel evacuation? We do not believe this interpretation to be adequate, because one method to chronically suppress singing in zebra finches for 4 or more days is to remain in close proximity to them and move a hand towards them as soon as they start to produce the first song introductory notes (we had used this manipulation in the past as a means to study the role of song practice on song learning). When we suppressed singing this way in three animals for up to 6 days, we did not observe any obvious aberrant behaviors to result from song suppression (unpublished findings). Therefore, there seems to be no reason to believe that singing is a vital need. The reason why none of our birds suppressed their songs for an entire week or longer as a result of isolation, surgery, tethering, and wearing an implant, we ascribe to the high coping capacity of zebra finches with regard to these manipulations.

It will be interesting to evaluate whether singing rate is also reduced or even suppressed by other types of stressors. A meta analysis of research data on zebra finches used as disease models might be particularly useful for evaluating the interplay between stress and singing rate. It will also be interesting to test whether in social conditions, stress resulting from visually confirmed conflicts or fights between animals is associated with reduced singing rate. Some evidence for this idea is provided by our data on bird pairs in which one bird sang a lot whereas the other hardly ever sang. We speculate that such extreme behavior signals a dominance relationship, in which the singer is more dominant.

It is currently not clear whether even small fluctuations in daily singing rate in a bird should be interpreted as concurrent waxing and waning of a small stressor. Some evidence against this idea is presented in Section 4.1, where a very mild stressor (white noise playback) does not affect singing rate, but nevertheless elicits an aversive behavioral response in most animals.

One obvious drawback of our measure is that it only applies to male birds (because females do not sing). We currently have no data from females that would allow us to test whether some of their call types could serve as a similar read-out for welfare.

## 3. Husbandry environment

### 3.1. Welfare associated with isolation and changes in husbandry

To understand certain behaviors and their neural mechanisms, it is important to perform experiments in controlled conditions. Individual housing (isolation) may be a necessary condition for some experiments. For example, isolate housing prevents damaging a neuronal implant. Also, isolation allows feeding a specific diet to individual animals (Reijmers et al., 2007). Acoustic isolation improves the quality of song recordings, because audio unmixing is not possible with multiple overlapping and highly variable sound sources such as birds (Pal et al., 2013). In zebra finches, single housing can also be an important behavioral parameter. Namely, non-directed (isolate) song is different from directed song (Bischof et al., 1981; Kao et al., 2005; Sossinka and Böhner, 1980), and the neural mechanisms of undirected (isolate) song are highly distinct (Hessler and Doupe, 1999; Jarvis et al., 1998). Thus, there may be scientific and practical benefits to isolate animals. However, what is the welfare impact of isolation? What is the value of transient presentation of known or unknown social partners to otherwise isolated animals? It is important to understand in what ways *single housing* and a *restrictive husbandry environment* impair animal welfare.

#### Stress due to isolation

Zebra finches are monogamous birds living in flocks of less than 10 birds to a few hundreds depending on the season (Zann, 1996) and therefore the default assumption is that they experience stress when isolated. However, the behavioral and physiological evidence for such stress is not that clear cut, even when compared with social housing conditions.

For one, there is a risk of harm associated with group housing. In the wild (Zann, 1996) and in captivity, (Bonoan et al., 2013; Ikebuchi and Okanoya, 2006), zebra finches form a dominance hierarchy displaying dominant and subordinate behaviors. In captivity, when zebra finch males are introduced into a new cage, aggression and fighting tend to occur. After a short time, a dominance hierarchy is formed and serious (physical) attacks are initiated mainly by the heavier and dominant aggressor against the lighter, subordinate defender. In a study, these roles of birds were stable and did not change over the course of an observation period of 10 days (Ikebuchi and Okanoya, 2006). In line with these findings, we have observed that sometimes birds stop singing in the presence of other birds (Fig. 2-6). Some male birds even modify the parameters of their songs when exposed to another male. Such observations indicate that companion birds or changes in group structure can be a source of behavioral adaptation (Gill et al., 2015; Vyssotski et al., 2016) and may even be a source of stress. It should be remarked that “in the wild, dominance hierarchies between individuals might not be established as clearly as in these experiment, because weaker males could escape from the site of stronger males before a firm hierarchy is established” (Ikebuchi and Okanoya, 2006). However, until further experiments are performed in more natural environments, we cannot disregard the possibility that intra-group dynamics can lead to distortions that could affect the conclusion of an experiment.

In some birds (e.g. parrots) there is risk of self-mutilation, feather plucking, and stereotyped behaviors, when they are held in environments lacking proper enrichment (van Hoek and ten Cate, 1998). In our lab, however, we have never seen zebra finches in isolation commit either self-mutilation or self-feather plucking, not a single bird out of more than 1000 birds has shown any signs of self-mutilation whatsoever.

A negative impact on welfare due to single housing is difficult to assess in zebra finches. Complete isolation in single cages (45 × 25 × 23 cm) causes an increase in corticosterone levels, but levels return to baseline after 30 minutes of isolation (Banerjee and Adkins-Regan, 2011). A similar study has shown an increase in corticosterone levels still after 12 hours of complete isolation in small cages (Perez et al., 2012). Thus, overall, the effect of complete isolation in small cages on corticosterone levels is not clear. Other results indicate that removal of vocal audience increases corticosterone levels in males, and alters temporal and spectral features of distance calls (Perez et al., 2012). In addition, as a social group increases in complexity, more new neurons are found in brain areas involved in vocal communication (Barnea et al., 2006; Lipkind et al., 2002). However, the effects of increased neurogenesis and neuronal survival on welfare are not known.

Male zebra finches isolated in cages of dimensions 28 × 30 × 39 cm perform all of the normal ranges of behaviors documented in birds (Slater and Ollason, 1972). Furthermore, completely isolated juveniles learn excellent song copies from song playbacks through a loudspeaker (Tchernichovski et al., 2001), revealing no learning impairment from isolation during development. Also, birds that grew up in isolation show afterwards no reproductive impairment (8/8 males that were raised in complete isolation in our lab have each bonded with a female and successfully raised their offspring).

Regarding stereotyped behaviors such as spot picking and route tracing, these would require a more detailed analysis. We have some anecdotal evidence that route tracing might happen (e.g., stereotyped jumping from one perch to another). However, at this moment, we cannot provide solid evidence due to lack of ethograms and absence of standardised records. With the newly devised automated beak tracking system (Section 2.1), it might be possible to analyse birds’ movements to detect stereotyped behaviors in a scientifically sound manner. So far, when we observed stereotypies indicating stress, we altered the environment inside the cage. In line with previous work, we have observed that changing the position of perches or adding a perch ameliorates or eliminates entirely the stereotypy (Keiper, 1969).

In Section 2.3 we argued that singing rate is an inverse measure of stress. If this measure is tenable, then our findings dramatically demonstrate that it is not isolation per se that is stressful to male zebra finches, but isolation in a new environment. Because in previous zebra finch studies examining stress from isolation, the confounding effects of environmental changes were either not controlled for or not accurately described, it remains to be determined which environmental changes are stress promoting upon isolation and which are not.

Important questions about isolation that will have to be addressed in future studies are the extent to which isolation and its resulting reductions of experiment duration and animal numbers are preferable for welfare compared to part-time or full-time social housing.

#### Stress from long-term isolation

We specifically inspected our data for evidence that long-term isolation is a source of stress. As illustrated in Fig. 2-6, we found that a long isolation period (following adaptation to the new environment) leads to a significant reduction in singing rate by on average 13 % (*range: −38 to +10%, p = 0.024, t-test, N = 6 birds)*. This finding might indicate that light stress could accumulate during chronic isolation. However, the drop in song rate mainly originated from a single bird that produced exceptionally many song motifs at the beginning of isolation (>4000 motifs, yellow curve in Fig. 2-6). At the end of the isolation period, this bird dropped its song rate to the range of rates in other birds. Thus, more birds will be needed to validate our observed effects of chronic isolation on singing rate.

Birds suffering from nutritional stress (malnourished birds) produce a song with reduced complexity, consisting of fewer syllable types per song motif compared to control birds (Spencer et al., 2003). We therefore tested whether birds, after being isolated for a long time, showed a degraded song and thus a possible sign of stress. In seven birds examined, we randomly picked 160 song motifs each both at the beginning and at the end of isolation (three birds were isolated for at least 28 days and four birds for at least 60 days). In four birds, there was no significant change in the number of song syllables per motif after long-term isolation (two-sample t-test, *p* >0.05 for each bird). In six birds, we found a small but significant increase in the number of syllables per motif (5.66→5.82: *t* = −3.2, *p* = 0.001; 5.53→5.74: *t* = −2.9, *p* = 0.004; 2.89→2.98: *t* = −2.1, *p* = 0.04; 4.93→5.24: *t* = −2.3, *p* = 0.02; 2.95→2.88: *t* = 2.5, *p* = 0.02, 4.18→4.63: t=-4.4, p=0.00001; two-sample t-test, *df* = 358 everywhere). Overall, these data demonstrate that there is no reduction in song complexity associated with long-term isolation (rather, isolation seems to produce a weak effect of increasing song complexity). Therefore, if song complexity were a reliable measure of past stress, then long-term isolated zebra finches are not stressed.

#### Stress from separation

Some experiments require partial social contact among animals. For example, to study the effects of social context on singing, males need to be presented with females to record their directed songs and they need to be separated from females to record undirected songs (Hessler and Doupe, 1999; Jarvis et al., 1998).

Instances of separation seem to induce stress. What we know so far is that separation of established zebra finch couples does not to lead to an increase in corticosterone (Crino et al., 2017; Schweitzer et al., 2014), although (Remage-Healey et al., 2003) found an increase after 24 hours. In Bengalese finches, repeated showing of a familiar female causes a decrease in directed singing (Toccalino et al., 2016), which could signal habituation or a reduction of interest.

What is the effect of frequent separation on song development in juvenile birds? Do juveniles continuously housed with a tutor learn their songs faster than juveniles raised with no more than 90 minutes of tutor exposure per day? To the contrary, we found that juveniles learned their songs faster when they were exposed to a male tutor for only 90 minutes per day. We traced the song learning in two groups of 4 juveniles each and subjected them to different tutoring paradigms. On 59 days post-hatching (dph), for each bird, we picked 20 pupil songs and 20 tutor songs to generate 400 comparisons. The average song similarity score of birds exposed only for 90 m/day and only for 14 days was higher than the average similarity score in birds that were continuously housed with their tutors for 24 days (Fig. 3-1; 49.2 ± 4.6 vs. 28.5 ± 5.4). We modeled song similarities as the linear mixture of fixed group effects (tutoring paradigm) and random individual (animal) effects (see Methods). A linear mixed effects model revealed the difference in song similarity score between the 24 h/day and the 90 min/day groups to be significant, with a fixed effect of 20.7 for the differences result only from tutoring paradigm (*t* = 2.41, *df* = 4797, *p* = 0.006; *N* = 4 birds in each tutoring group). Faster learning by juveniles with limited exposure was not due to the social isolation: when we housed 4 additional juveniles continuously with an adult female for 24 days, then the same 90 m/day of tutor exposure during 24 days resulted in a high average song similarity score of 46.5 ± 7.9. This latter group was associated with a fixed effect of 18.0 in excess of the 24 h/day group (*t* = 2.41, *df* = 4794, *p* = 0.016).

**Figure 3-1.**
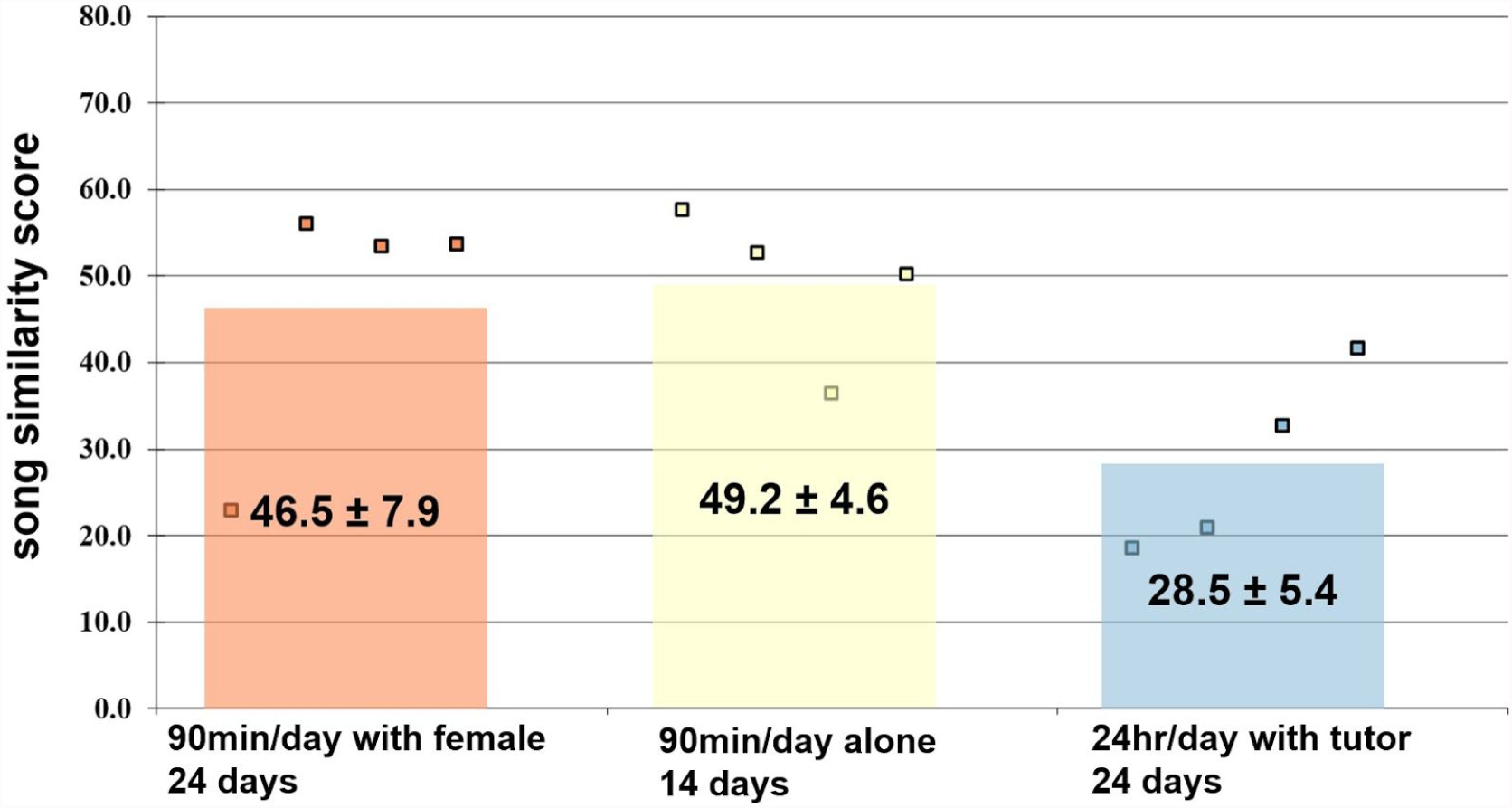
Juveniles with restricted exposure to a male tutor learn faster than juveniles that are housed with their tutor. In three groups of juvenile males we assessed the speed of song learning using the song similarity score (SAP Pro) between the juvenile’s song evaluated at 59 dph and the tutor’s song. The average song similarity score in juveniles that spent 90 minutes per day with their tutor (orange bar, yellow bar, N = 4 birds each) was higher on 59 dph than the average score in four juveniles that spent 24 h per day with their tutor (blue bar). Birds in the 24 h/day group (blue) were housed for 24 days with their tutor until 59 dph. Birds in the 90 min/day group were either housed for 24 days with a female (orange), or for 14 days in isolation (yellow). The colored squares indicate average song similarity scores in individual birds.

Thus, juveniles routinely separated from their tutors learn faster than juveniles with unlimited tutor exposure, regardless of the husbandry condition (isolate vs. social housing). Overall, these findings agree with findings in playback experiments, where it was found that less exposure to tutor song leads to better song copying (Tchernichovski et al., 1999). We conclude that there is stress from separation and isolate housing that would negatively impact song development.

#### Body weight

Here we investigate whether changes in husbandry condition can affect body weight. As part of a general health screening, we perform body weight measurements to determine how individual birds cope with their new situation. We gathered health-monitoring data associated with changes in housing and social conditions and inspected the data for evidence of loss of body weight.

We have found that adult birds have a tendency to lose weight in the days following a change in husbandry (Fig. 3-2B). Birds lost weight when they were removed from a social housing condition and isolated in a new environment (*N* = 3 birds, 4.8%, 5.7%, 2.9% weight loss). Also, when we exchanged two adult males in a group of four adult males, we found that the newly arriving birds tended to lose weight (7.0% and 5.2% weight loss), and so did the two remaining birds (5.4% and 6.7% weight loss). When isolated birds were transferred to a social group, two of three birds maintained their weight (+0.19% and −1.9% weight change), whereas one bird lost weight (6.5% weight loss), raising the possibility that the transfer to a social condition is less stressful than isolation in a new environment (Fig. 3-2B). However, data from more animals are needed to corroborate this possibility.

**Figure 3-2.**
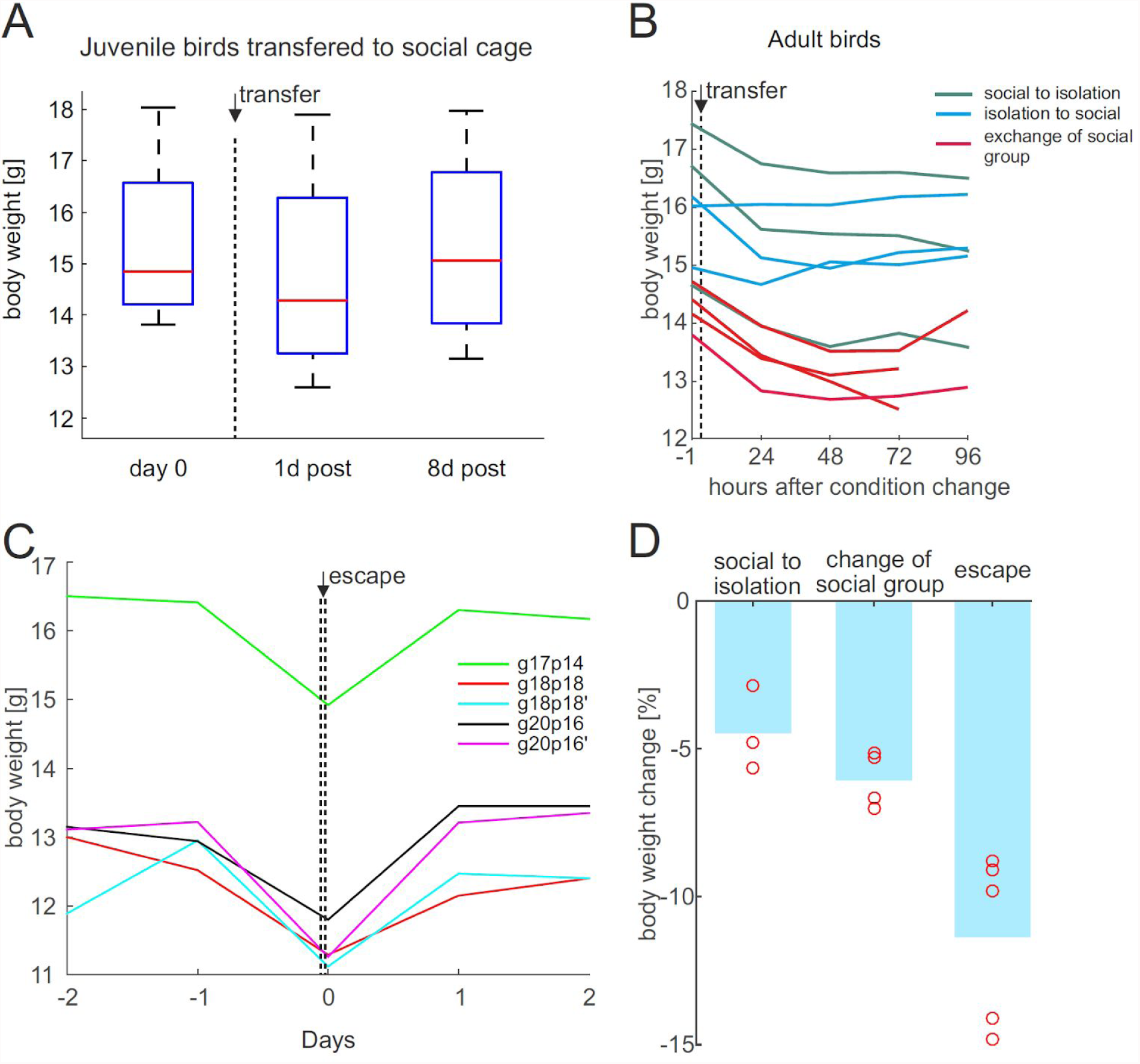
Weight loss due to change in housing condition is smaller than weight loss during a brief 5-minute escape. (A) Juvenile birds (age 76-102 days) lost weight after separation from parents and transfer to a social cage. The body weight was regained after 1 week in the social cage. (B) Birds lost weight upon transfer from a social cage to isolation in a new environment (green lines, ‘social to isolation’, relative weight loss also shown in D). Four birds lost weight upon exchange of social group (red lines, two birds were moved to another social cage and the two remaining birds received new companions in their cage, see also ‘change of social group’ in D). Only one of three birds lost weight upon transfer from isolation to a social cage (blue lines). (C) Body weight decreased during a brief < 5-minute flight in the laboratory room upon escape from the social cage. Body weight returned to pre-escape levels in 1–2 days. (D) The average weight loss for the escaped birds in C was −11.3% (’escape’), much larger than the weight loss upon cage transfer (‘social to isolation’, and ‘change of social group’).

We also inspected body weight in juveniles that were transferred away from their parents to single-sex aviaries containing a maximum of 19 birds at 76–102 dph. The juveniles lost weight within one day (*N* = 4 birds, 8.8%, 4.8%, 2.8%, 0.7% weight loss respectively), but they tended to regain the weight within one week after the transfer (Fig. 3-2A).

In a few cases, by chance we measured body weight before and after an instance of escape (Fig. 3-2C). That is, birds occasionally escape from a cage while an animal caretaker or experimenter tries to capture it. We usually wait for escapees to calm down and perch before we approach them with a net. Surprisingly, when we measured body weight immediately after such brief escapes lasting no more than 5 minutes, we found an average weight loss of −11.3% relative to the preceding day (Fig. 3-2D). The weight loss might have been caused by evacuation of liquids (e.g. in feces). The weight loss was transient, as birds regained their weight within 1–2 days. Thus, short flights outside the cage can lead to a remarkable loss of body weight within just a few minutes.

In summary, we find that adult zebra finches tend to lose weight when isolated from a social group or when a conspecific is exchanged from a group. However, the loss of body weight we measured is small, about 2–3 times smaller than the typical loss resulting from a brief <5-minute escape from the home cage. Overall, these findings on body weight point to a non-substantial burden associated with changes in social condition.

Given these data, what are the tradeoffs between isolate housing and partial social housing? Overall, it is difficult to assert whether stress from temporal separation (partial social housing) is worse than possible chronic stress from isolate housing, for one because there is no unambiguous evidence for the latter. Also, the welfare benefit of part-time social housing remains questionable: whereas some data showed that separating birds tends to be stressful for both parties, this stress might be counterbalanced by the reward of their reunion. However, there is currently very little data to support this idea.

### 3.2. Enrichment with mirror

Can mirrors provide benefits to birds in captivity?

Only a handful of animals have passed a mirror self-recognition test (Gallup, 1970) without any training (Suddendorf and Butler, 2013). Most animals, “when initially exposed to mirrors, respond as if the image represented another animal” (Gallup, 1970). Indeed, many studies show that mirrors can mitigate the effects of social isolation in animals including birds (Henry et al., 2008), cattle (Piller et al., 1999), horses (Kay and Hall, 2009), rabbits (Dalle Zotte et al., 2009), mice (Watanabe, 2016), sheep (Parrott et al., 1988), rhesus monkeys (Gallup and Suarez, 1991), and stumptail monkeys (Anderson, 1983); although some studies found no effect in rabbits (Jones and Phillips, 2005) and mice (Fuss et al., 2013; Sherwin, 2004). In rats, mirrors have little effect (Reed et al., 1996).

In horses, for example, the use of mirrors reduces the display of stereotypical behaviors (McAfee et al., 2002). It is possible that the effect of a mirror as a surrogate companion disappears over time due to habituation. However, reinstating the mirror after a break of five days causes monkeys to react again with social responses towards it (Gallup and Suarez, 1991).

In many bird species, including kea, starlings and domestic fowl chicks, mirrors reduce the isolation stress in laboratory conditions (Diamond and Bond, 1989; Henry et al., 2008; Montevecchi and Noel, 1978). For this reason, in order to mitigate the possible burden of social isolation, we add a mirror outside the cage of isolated birds. Contrary to territorial birds or birds with strict dominance hierarchy (Sumitani, 2014), the use of mirrors in social birds mitigates the effects of limited social contact (Henry et al., 2008; Keiper, 1970). Furthermore, zebra finches develop a preference towards a mirror compared to a conspecific (Gallup and Capper, 1970; Ryan, 1978), and the same effect was also seen in rabbits (Zotte et al., 2009).

Here we present preliminary data from an experiment in which we repositioned a mirror, suggesting that mirror repositioning can constitute an enrichment that prevents boredom and possibly suppresses stereotyped movements. Using our automatic beak tracking system (Section 2.1), we found that a bird changed its location preferences in the cage when we repositioned the mirror (Fig. 3-3). The spatial position map of the bird was very different before and after mirror repositioning.

**Figure 3-3.**
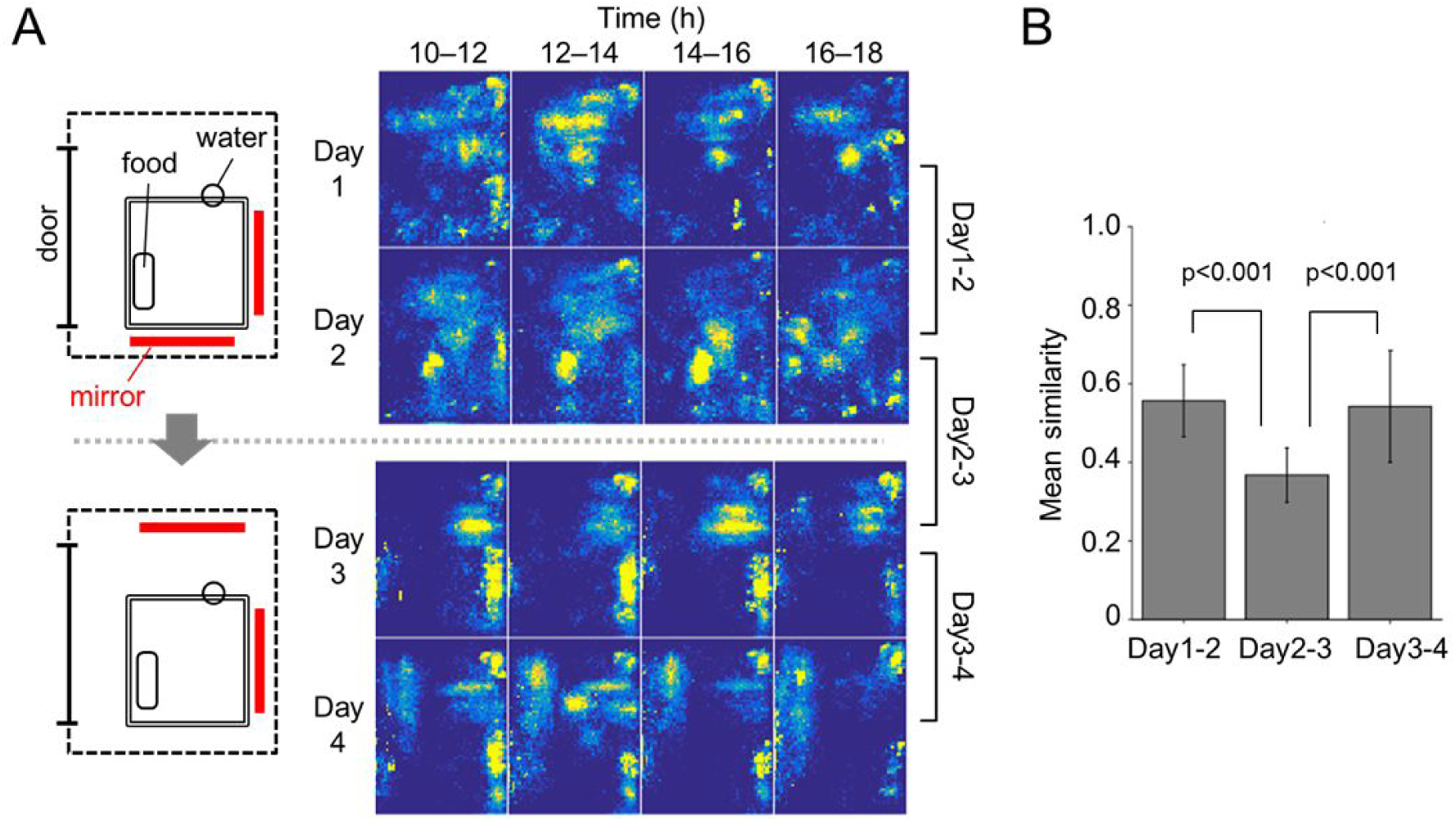
Mirror repositioning induces new place preferences. (A) The position maps in 2 h windows across two days before mirror repositioning (top) and after mirror repositioning (bottom). The mirrors are indicated as red bars in the top view of the bird’s cage (right). (B) Mean similarity of position and movement maps across consecutive days is high on days before (Day1-2) and after (Day3-4) mirror repositioning, but small (Day2-3) across the mirror repositioning event. The similarity was calculated as the Pearson correlation coefficient of time-matched maps across consecutive days. Significant differences were observed between Day1-2 and Day2-3 (two-sample t test with Bonferroni correction, t(30) = 6.59, p < 0.001) and between Day2-3 and Day3-4 (t(30) = −4.42, p < 0.001).

We have also noted that isolates with mirrors tend to be less frightened by an approaching experimenter than isolates without mirror. This same observation has been made in other laboratories as well (D. Lipkind, personal communication). Presumably, the mirror mediates adapted behavioral responses to moving stimuli. Thus, we suggest that the use of mirrors as social substitutions in isolated animals can mitigate the effects of isolation by diminishing aberrant behaviors and reducing stress, and a routine change of the mirror location can increase animal’s mobility and might help to prevent stereotyped behaviors.

### 3.3. Interactive auditory and visual communication system

Can an interactive telecommunication system in isolated animals restore some of the natural vocal exchanges between animals?

Video and audio playback are potent techniques to study animal communication. Methods such as interactive playback and virtual social environments (that simulate social environments) have been successfully used to study social interactions and social influences during developmental learning (King, 2015; Ljubičić et al., 2016). As much as such interactive playback systems (IPS) have helped to advance our understanding of the roles of communication signals in many animal taxa, we argue that further advances will result from interactive communication systems (ICS) that allow for controlled vocal communication between two or more animals.

#### Interactive playback systems (IPS)

Video stimuli have been shown to elicit natural behaviours in some animals: chimpanzees and budgerigars yawn more in response to videos of conspecifics yawning than to control videos in which conspecifics were engaged in other behaviours (Campbell and de Waal, 2011; Gallup et al., 2015); gloomy octopus readily approached and touched the video screen when presented with a crab video, and reduced their activity, a natural response for this solitary species, in response to a video of a conspecific (Pronk et al., 2010); videos of ‘audience’ hens (*Gallus domesticus*) potentiate alarm calls produced in the presence of a predator model (Evans and Marler, 1991) and male Jacky dragons (*Amphibolurus muricatus*) produce aggressive displays in response to videos of conspecific males (Ord et al., 2002); budgerigars copy the actions of a live or video demonstrator in a two-action test (Heyes and Saggerson, 2002; Mottley and Heyes, 2003) and Burmese red jungle fowl (*Gallus gallus spadecius*) copy foraging choices of live and video demonstrators (McQuoid and Galef, 1992, 1993). In some cases, there is no qualitative difference in the response to live versus video stimuli (Evans and Marler, 1991; McQuoid and Galef, 1993; Ord et al., 2002; Rieucau and Giraldeau, 2009), however, in other cases an attenuated (Ikebuchi and Okanoya, 1999) or enhanced response to video stimuli was reported (Swaddle et al., 2006), suggesting that interactions among animals may exhibit dynamics that are hard to mimic using playback systems.

More specifically, male zebra finches and Bengalese finches sing directed song to video presentations of female conspecifics (Ikebuchi and Okanoya, 1999) and female zebra finches will perform courtship display to videos of male conspecifics (Swaddle et al., 2006). Zebra finches also learn social information about foraging from video demonstrators (Guillette and Healy, 2014, 2017). Live streaming has a similar effect on observers’ behaviour as live demonstration (Guillette and Healy, 2017). When looking at developmental learning, juvenile zebra finches learn much better from real tutors than from tutoring using interactive playback (Derégnaucourt et al., 2013).

#### Interactive communication system (ICS)

Some of the limitations of playback systems suggest that to examine the function of vocal and visual interactions, it may be important to study exchanges not between a bird and a robot or computer, but between two or more birds. As a first step, we have developed a custom digital telecommunication system that allows for visual and auditory contact between birds. Our system allows us to control the audio and visual channels independently. In uses of that system, we have observed that adult zebra finches engage in reliable vocal interactions (Fig. 3-4). We have also observed that juveniles interact with adult males in that system and they are able to learn good song copies from them. One of the important future uses of this system will be to digitally manipulate vocal interactions between animals in order to identify the vocal signatures that influence song learning. For example, a key application could be to substitute a call type by another one to detect the roles played by these calls during learning.

**Figure 3-4.**
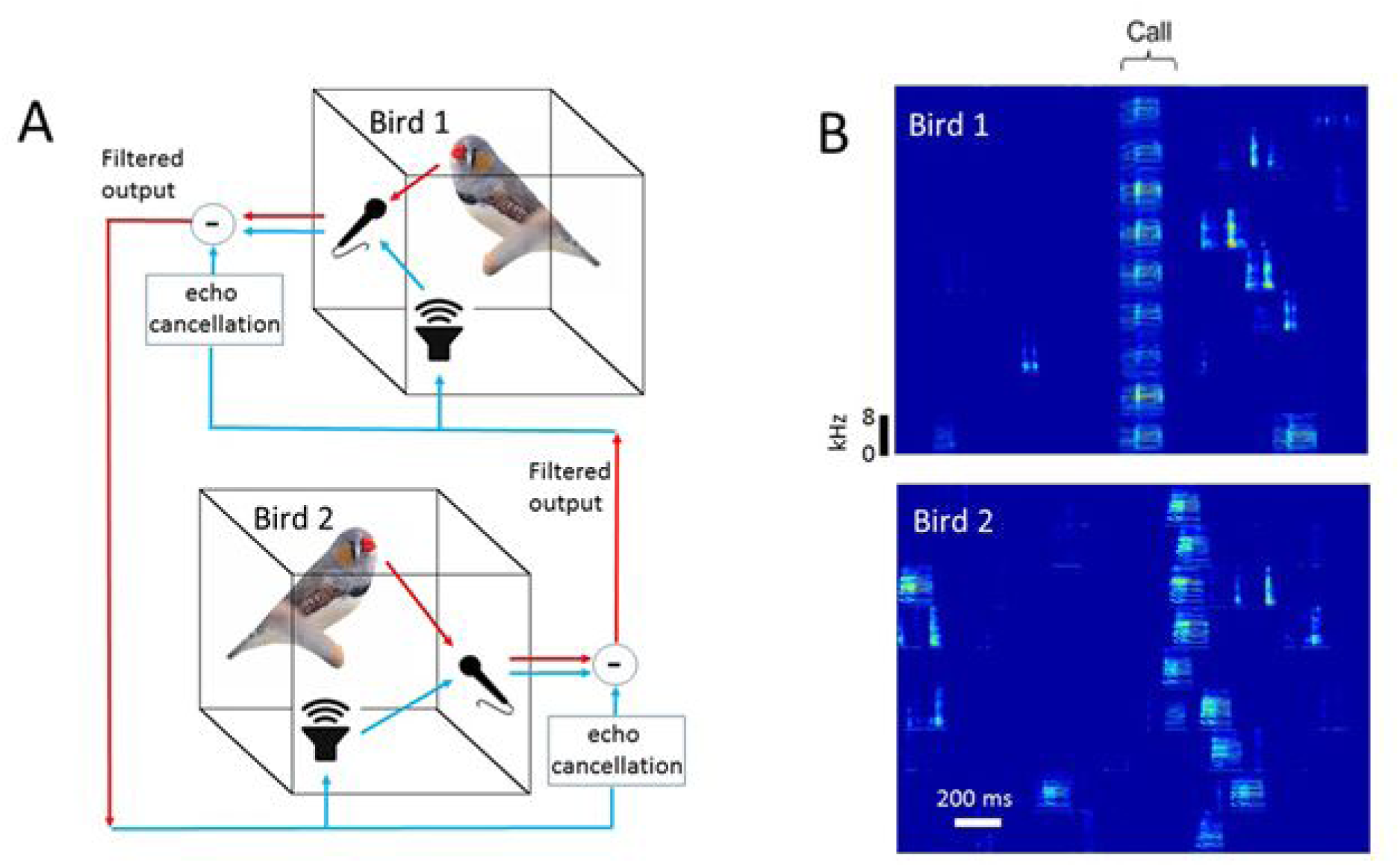
An interactive auditory and visual communication system promotes robust call interactions. A) Birds are kept in soundproof boxes. They are in visual contact either via two windows or a pair of cameras and computer screens. They are in auditory contact through a pair of microphones and loudspeakers endowed with echo suppression. The echo suppression system subtracts the sounds played through the loudspeaker (blue) in real time from the microphone signals, to result in a filtered signal that constitutes a relatively clean recording of the sounds generated by the bird (red). B) Nine example calls produced in one male bird (top). In 90 minutes, the bird produced the shown call type 203 times, 38 out of which were answered by a call (bottom) in the other male within a 500 ms window following the onset of the first call. Thus, this type of call-call response occurred 10 times more often than by chance (20% versus a chance level of only 2%), demonstrating naturalistic call interactions (Anisimov et al., 2014) in this bird pair.

We expect that the possible effects on animal welfare of our system will strongly depend on the manipulated signal. For example, whereas the suppression of distress calls may reduce stress, the introduction of such calls into vocal interactions could increase stress.

### 4. Behavioral and peripheral manipulation

### 4.1. Song conditioning

What is the impact of song operant conditioning on welfare?

To reinforce certain actions is one of the most intuitive ways of teaching: for example parents use reinforcement to teach their children when they smile at them or when they scold them. Operant conditioning has become famous with Pavlov’s Nobel Prize in 1904 and is one of the most widely applied strategies used to train animals (Hollerman et al., 1998; Kravitz et al., 2012; Montague et al., 2004; Rueda-Orozco and Robbe, 2015; Tumer and Brainard, 2007; Wolpert et al., 2011) and humans (Galea et al., 2015; Montague et al., 2004; Nikooyan and Ahmed, 2015; Schönberg et al., 2007; Shmuelof et al., 2012; Wolpert et al., 2011).

Zebra finch song can be operantly conditioned using loud acoustic stimuli as aversive reinforcers (Ali et al., 2013; Andalman and Fee, 2009; Charlesworth et al., 2012; Tumer and Brainard, 2007). Song conditioning has been used to show that the spectral tuning of song is actively controlled by the basal ganglia (Ali et al., 2013; Andalman and Fee, 2009; Charlesworth et al., 2012; Tumer and Brainard, 2007). Moreover, when the aversive reinforcer is withdrawn in such experiments, birds gradually retrieve their original songs they produced before the aversive reinforcer was introduced. Higher brain areas selectively involved in this song retrieval process have been identified (Canopoli et al., 2014). Thus, by combining behavioral conditioning and brain perturbations, brain areas can be identified that are discriminatively involved in either adaptation to environmental influences or in memory retrieval (Ali et al., 2013; Canopoli et al., 2014).

There is a high potential benefit of such studies to humans. A better understanding of the interplay between motor exploration and reward within the architecture of the vertebrate brain could serve to improve treatment of brain disorders. For example, principles of adaptive motor plasticity could be applied to design better rehabilitation strategies in patients with neurologic disorders. Unfortunately, although patients can quickly normalize both reaching movements and gait in adaptation paradigms, abnormal movements tend to quickly return once the perturbation is removed (Reisman et al., 2007; Scheidt and Stoeckmann, 2007). It was recently shown that binary feedback as used during song conditioning in zebra finches and Bengalese finches can drastically increase the amount of learning retained in human adaptation paradigms (Shmuelof et al., 2012).

Noise, including white noise from streams and waterfalls, is part of the natural environment, but also in captive environments there is frequent noise. Songbirds can deal with environmental noise, for example by changing the pitch and loudness of their song (Dominoni et al., 2016; Luther and Baptista, 2010; Slabbekoorn and Halfwerk, 2009). This trait is exploited in laboratory experiments in which playback of white noise is used as an aversive stimulus. To our knowledge, the burden of song conditioning by means of acoustic stimulation has not been studied in detail.

In our past work on song conditioning, we have monitored birds’ behavioral responses to brief (50 ms) but loud white noise stimuli delivered at precise times within the song motif. To avoid damage of hair cells in the zebra finch cochlea (the basilar papilla) (Ryals et al., 1999), we limited sound pressure levels of white noise stimuli to 100 dB. Typically, we broadcast white-noise stimuli from a loudspeaker at a sound pressure level corresponding to 85 dB at the bird’s ear. This pressure level is within the natural range of the bird’s own song. Namely, song syllables can reach up to 85 dB at a distance of 25 cm above bird’s head (Ritschard and Brumm, 2011). We delivered white noise contingent on a certain sound feature within a specific target syllable, such as for example the pitch (fundamental frequency) of that syllable or its duration. Typically thresholds for white noise delivery were such that about 50% of target syllables were hit by white noise, and 50% were not.

Assuming that the white noise stimulus is aversive to birds, they could try to avoid this aversive stimulus either by singing less or by suppressing the relevant sound feature. We found no evidence for the former. Birds did not decrease production of the target syllable within the three days following onset of song conditioning, compared to the three days just before the onset (*N* = 14 adult male zebra finches; t-test, *p* = 0.57; see Fig. 4-1). Also, no decrease in syllable production rate was observed on the first day of playback onset compared to the day before (*N* = 14 adult male zebra finches; t-test, *p* = 0.88).

**Figure 4-1.**
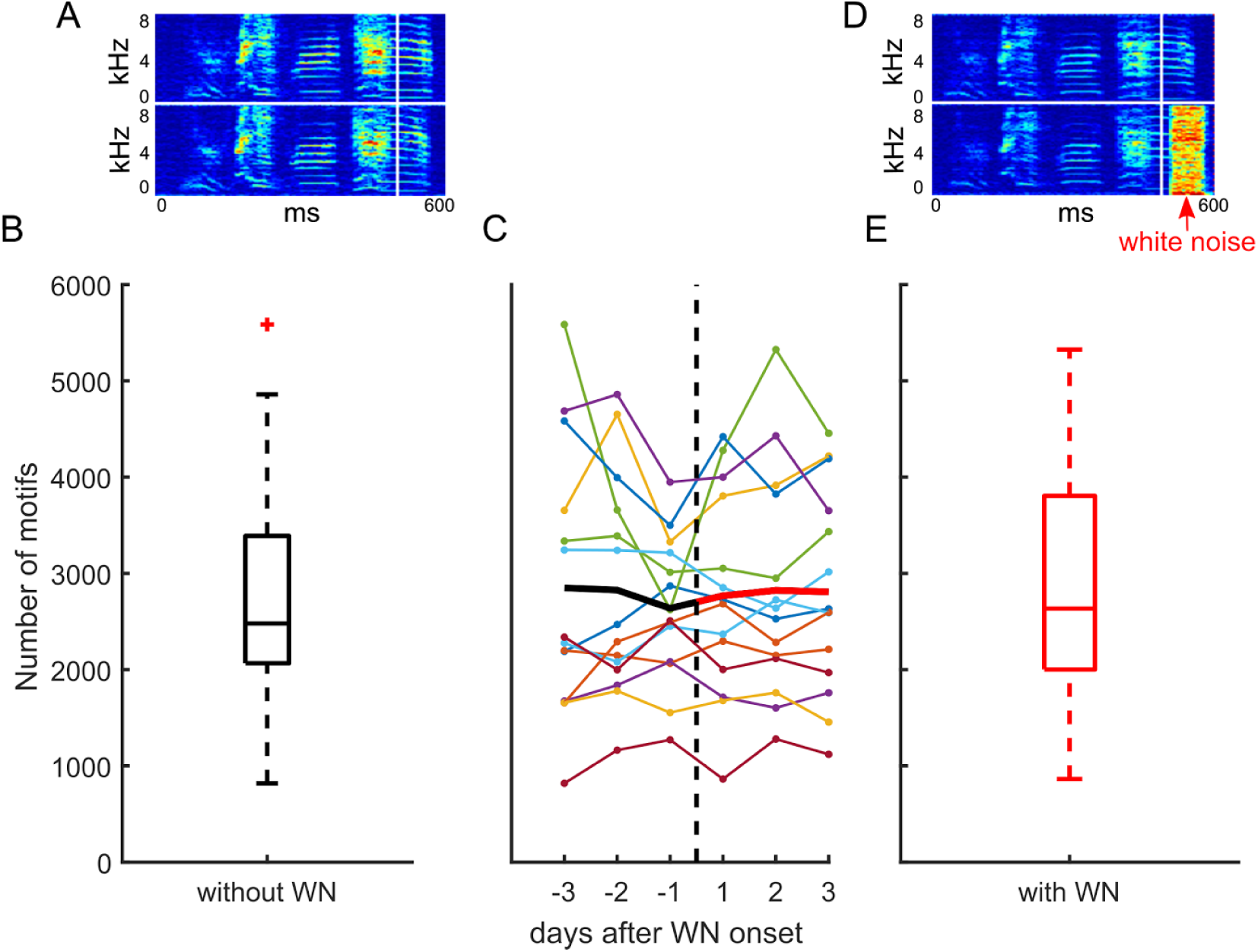
Singing rate in adults is unchanged by onset of interactive white-noise playback. A. Example spectrograms of unperturbed song. B. Box plot of the number of song motifs produced per day during the last three days before onset of interactive playback. C. Number of motifs per day in each bird (colored lines) as a function of the days before and after onset of interactive playback. The black line shows the average across birds before playback onset and the red line the average after playback onset. D. Example spectrogram with and without white noise. E. Box plot of the number of motifs per day during the three first days of interactive playback. Playback of white noise (WN) was triggered whenever the pitch of the targeted syllable was below (or above) a manually set threshold. Birds could avoid white noise playback by stopping to sing. However, the average number of songs produced before and just after onset of playback (within three days each) were not different from each other (N = 14 adult male zebra finches).

Nevertheless, most birds (27/38 birds) responded aversively to white noise, because they changed the sound feature of the targeted syllable in such a way that fewer syllable renditions triggered white noise (these birds shifted the syllable feature away from the white-noise triggering region). Some of the remaining birds (9/38 birds) did not change the sound feature of the targeted syllable, as if they did not care. Most interestingly, a few birds responded by singing more syllable renditions that triggered white noise (2/38 birds, Fig. 4-2). That is, in these two birds, the percentage of target syllables triggering white noise playback was significantly higher in the evening compared to the morning. In the morning, during the first 200 song motifs, 55% ± 14% of target syllables triggered white noise whereas in the evening, during the last 200 daily song motifs, this number was 76% ± 15% (t-test, *p* < 0.001; 25 days). One bird received white noise when a particular song syllable was shorter than some daily fixed threshold, the other received white noise when the pitch of a particular syllable was below threshold. Both birds seem to have been attracted by interactive white noise, not repelled by it. In these birds, playback of white noise stimuli during singing seemed to represent an enrichment rather than a burden.

**Figure 4-2.**
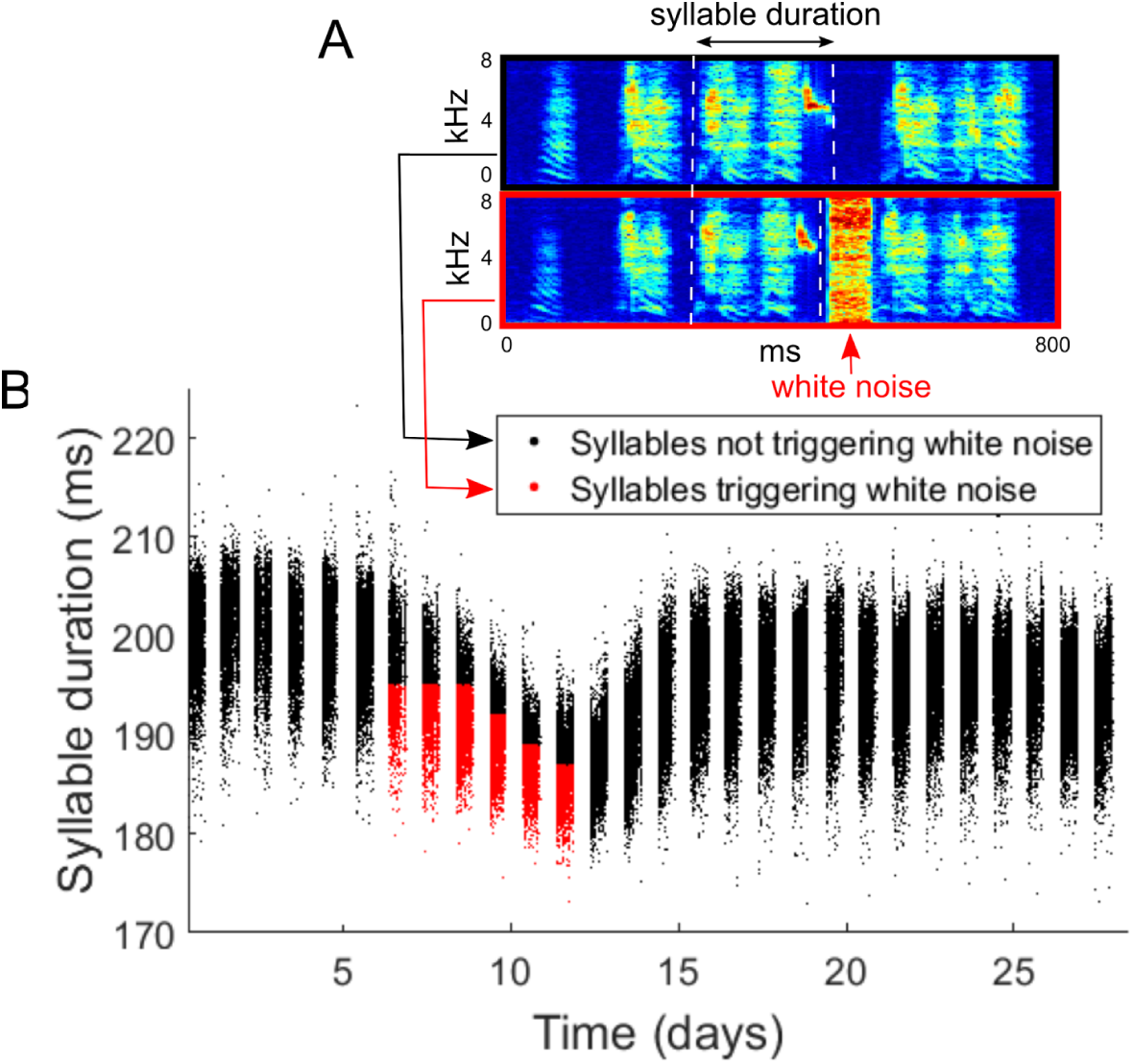
Example bird that shifts the targeted acoustic song feature (syllable duration) towards the white-noise triggering region. A) Example spectrograms of a song motif that did not trigger white noise playback (black box) and a motif that triggered white noise playback (red box). The white dashed lines mark the onsets and offsets of the target syllable. B) Scatter plot of target syllable duration as a function of time. Black dots mark long syllable renditions outside the white-noise triggering zone, and red dots mark short renditions that triggered white noise playback. This bird time-compressed the targeted syllable and thus modified its song to receive white noise more frequently.

Overall, we find that isolated birds do not reduce their singing when white noise playback starts, suggesting that this manipulation is associated with little burden. A mild burden is signalled in most animals because they adapt their song to avoid white noise. In other animals, however, white noise can serve the function of environmental enrichment, following the rule that aversive situations trigger stress responses only if the individual perceives them as aversive. We speculate that the burden of acoustic stimuli used in experiments strongly depends on sound intensity levels, implying that there might be a range of sound intensities in which interactive stimulus playback is experienced as enrichment in most animals. This speculation would be in line with practices in zoological gardens, where sensory stimulation is used to reduce pathological behaviors caused by “boredom”, inciting animals to increasingly perform their natural behaviors (Martínez-Macipe et al., 2015; Wells, 2009).

### 4.2. Muting by air sac cannulation

What do manipulations of the air sac mean for the welfare of zebra finches?

Birds can be muted by opening the air sac in their respiratory system. One of the main benefits of muting is to probe the effects of singing on a given behavior or on social status. By muting wild territorial songbird males and returning them to their natural environment, the role of vocal communication in territorial defence and in reproductive success have been discovered in Scott’s seaside sparrows (McDonald, 1989), in red-winged blackbirds (Peek, 1972), and in red-winged blackbirds (Smith, 1979). When the muting was reversible, the long-term effect of this manipulation was minor given that upon voice recovery, the birds were able to mate and acquire territory (McDonald, 1989).

The roles of singing (practice or rehearsal) in song learning and in song maintenance are not completely understood, which is why muting remains an important manipulation for addressing such crucial questions. For example, currently, it is not known whether zebra finches can retrieve any song from memory within just a few trials, or whether for some songs they need to practice several hundred times before retrieving these songs. This question is relevant because the vocal abilities of birds may constrain their cultural learning and vocal communication behaviors. Below we argue muting by air sac manipulation is currently the most suitable manipulation for suppressing song for a selected period of time.

A minimally invasive procedure to mute an animal is air sac cannulation. Air sacs are made of thin but strong membranes that are easily accessible by a small incision into the skin. Insertion and fixation of a cannula are relatively simple and quick procedures (Brown and Pilny, 2006; Graham, 2004).

Air sac cannulae prevent the buildup of air pressure to levels sufficient for vocalization. Based on our observations, animals with air sac bypass can still produce calls, but longer vocal sequences (e.g. songs) are partly or fully suppressed. When air sac cannulae are equipped with a valve that can be precisely controlled, selected song notes can be manipulated in terms of their acoustic properties, allowing the study of biophysical sound generation mechanisms (Amador and Margoliash, 2013).

Cannulation is a commonly used procedure to treat pathologies of the air sac system. It is applied in case of respiratory tract obturation and for ventilation (Brown and Pilny, 2006). In addition, cannulation is a convenient procedure for delivering isoflurane in zebra finches during surgery when access to the head is required (Nilson et al., 2005). Cannulae can be left implanted for several days (Brown and Pilny, 2006).

The effect of cannulation on the welfare zebra finches has not been studied in detail. A study on ducks has shown that cannulation affects two physiological parameters (Rode et al., 1990). That is, tidal volume and minute ventilation were significantly higher after air sac cannulation (160% and 200% of baseline values, respectively). It is not clear whether higher tidal volume and minute ventilation can be a cause for secondary negative effects, because avian lungs are not elastic and the pressure in the alveoli does not change during breathing as it does in mammals. The following parameters were not affected by cannulation: PaCO_2_, heart rate, and mean arterial blood pressure. None of the eight ducks developed any symptoms of respiratory tract disease. Post mortem analysis of the ducks’ air sac systems showed in one duck fibrinonecrotic air sacculitis due to aspergillus infection. It is not clear whether the infection was already present before the experiment. To our knowledge, none of the following symptoms of respiratory tract diseases listed in (Graham, 2004) have been reported in scientific publications on air sac cannulation in healthy animals: sneezing, yawning, scratching of the nose or oral cavity, open-mouth breathing, tail bobbing, apparent sternal movements, gurgling on inspiration or expiration, and head shaking. Overall, we estimate the burden from cannulation and muting to be small and mainly signalled by the small increase in respiratory rate.

In our view, muting via cannulation is less invasive and, less burdensome than other muting procedures, four of which are discussed in the following. 1) Muting can be performed by implantation of immobilizing pins into the syringeal cartilages (Cooper and Goller, 2004). This approach requires rupturing of the interclavicular air sac to gain access to the syrinx, so it is at least as invasive as simple cannulation. Similar to cannulation, this muting procedure is reversible, but unmuting requires a second surgery for pin removal (in cannulation, no second surgery is required, the animal is unmuted by insertion of a plug into the cannula). 2) Muting can be performed by cutting or crashing the tracheosyringeal nerves (TS) to denervate the syringeal muscles (Liberti et al., 2016; Peek, 1972; Solis et al., 2000; Tang and Wade, 2014; Vicario, 1991). However, this manipulation tends to cause severe respiratory problems including choking, which is why denervation might be more burdensome for the animal than cannulation (Smith, 1979). The TS nerve can be cut unilaterally to mute certain notes in birds with specialized laterality of the syrinx (for example in canaries but not zebra finches), or in zebra finches, to disrupt the spectral organisation of song syllables (Liberti et al., 2016). This latter procedure seems to be tolerable to the respiratory system, but complete denervation is not reversible, so it may be unsuitable to address certain types of questions. TS nerves can regenerate after nerve crush but song restoration is not complete and the recovery takes at least 4 weeks (Bhama et al., 2011). 3) Tracheostomy can be performed to mute a bird. However, the sound source (syrinx) in birds is below the trachea and not above it, as the vocal folds in mammals. Therefore, making an opening in the trachea tends to result in filtering of the produced sounds, but not in muting. In the past, we have performed tracheostomies in zebra finches to attempt to mute them, but 6/6 birds could vocalize after the surgery, often producing rhythmic songs with disrupted harmonic structure. Further problems may occur with tracheostomies such as swelling of the tracheal tissues or in case a foreign body enters the trachea. In a study that describes usage of chronic tracheostomies in parrots, sudden death of the animals was reported after 3-4 months due to problems with the respiratory system (Lennox and Nemetz, 2005). 4) A classical technique to de-vocalize noisy birds is to perform a surgery that exposes subsyringeal and suprasyringeal airways to the same respiratory pressure within the interclavicular air sac, presumably eliminating the pressure difference and thus the air flow through the syrinx. In this surgery, the syrinx is first exposed through an incision in the interclavicular air sac. Then, an opening is made in the trachea rostral to the syrinx, continuing midsagittally between the paired ventralis muscles (Phan et al., 2006). “After this procedure, males engaged in song-like beak movements and did not phonate, but onrare occasions produced short on rare occasions produced short breathy sounds” (Phan et al., 2006). Such devocalization procedure is irreversible and seems to induce a comparable burden as method 1 above using an immobilizing pin.

In our experiments, implantation of a cannula into the abdominal air sac strongly mutes zebra finches’ songs for several days. In these experiments, we inspect the cannula daily. If mucus was present or airflow through the cannula was reduced, we cleaned the cannula with sterile tissue and saline (airflow through the cannula was verified by holding in front of the opening a small feather). We monitored the bird’s vocal activity using our real-time song detection system. If a song was present, the cannula was inspected and cleaned.

We performed our first experiments using polyethylene cannulas (diameter 1.5 mm). Four birds were fully muted after cannula implantation for a duration of 5 days, 2 days, 1 day, and 2.5 days, respectively. Following these periods of complete muting, the birds produced a few song motifs daily (at 2–12% of their normal song production rates). With time it became more difficult to fully clean the implanted cannulae because the cannula ending inside the air sac tended to get clogged. The cannulae remained closed and birds were unmuted on days 7, 6, 6, and 4, respectively. Once the cannulas were closed, the unmuted songs became indistinguishable from the pre-surgery songs (Fig. 4-3).

**Figure 4-3.**
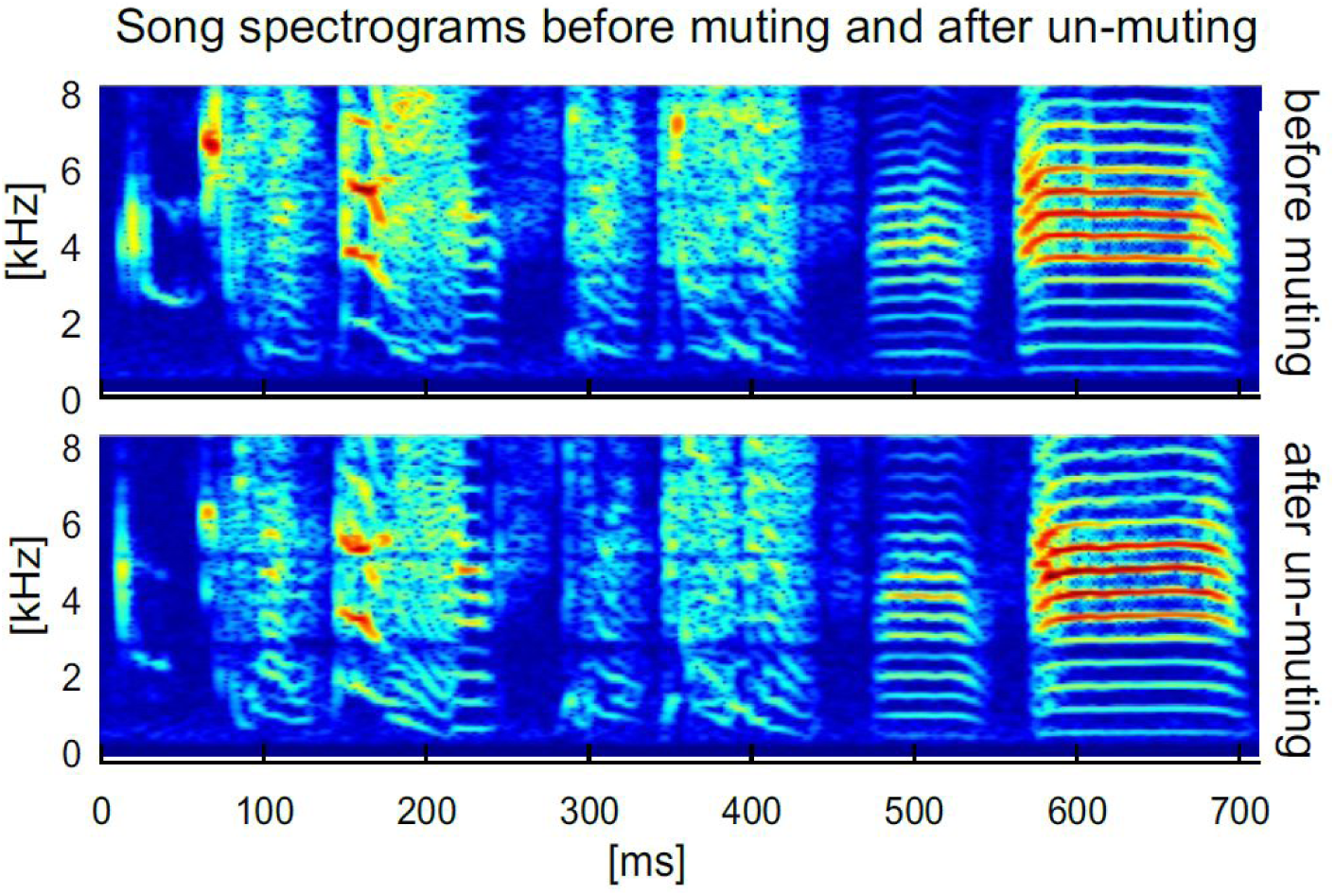
Zebra finch song before muting and after unmuting is identical. The song motif before the muting procedure (top) is visually indistinguishable from the song recorded after unmuting (bottom). Shown are logarithmic sound spectrograms of an example bird.

**Figure 4-4.**
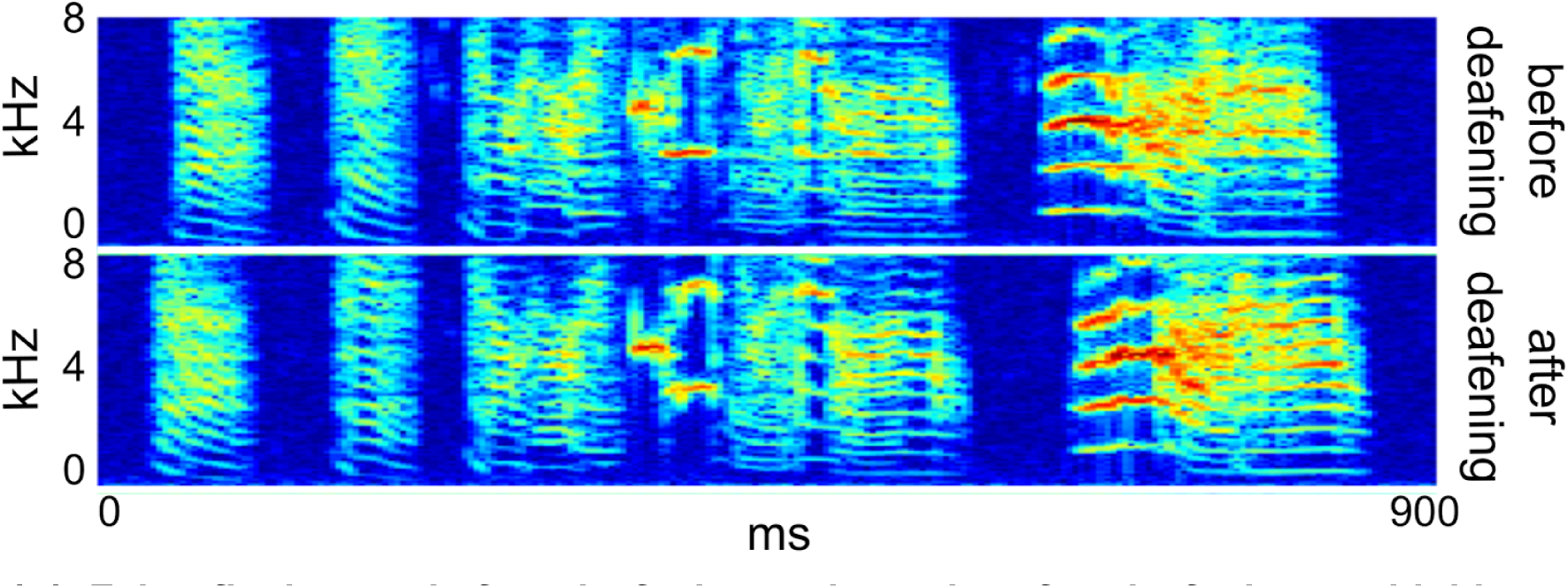
Zebra finch song before deafening and one day after deafening are highly similar. Shown are logarithmic sound spectrograms of an example bird produced one day before (top) and after deafening (bottom).

The number of song motifs produced in the unmuted state was identical to the pre-surgery singing rate. In three birds, the 6-day average number of songs motifs produced before cannulation were identical to the 6-day average following unmuting (two-sample t-test, *p* = 0.87, *p* = 0.68, *p* = 0.12, 6 days before and after muting); in the fourth bird, the number of song motifs produced after unmuting was slightly lower than before cannulation (average number of song motifs per day before muting: 1825 ± 300 and after unmuting: 1293 ± 78; *p* = 0.01). The preserved singing rate in the unmuted state suggests that cannulation does not produce a substantial discomfort during song production.

Two subsequently used birds were implanted with polyimide cannulas (diameter 1.5 mm). This material reliably prevented mucus formation and birds were fully muted for 8 days.

Unmuting in these animals also occurred spontaneously after 9 days. Overall, these preliminary data suggest that polyimide is highly preferable over polyethylene as tubing material for cannulation, because the resulting muting is more complete and the need for handling is minimized. The burden of cannulation seems minimal but could be better estimated in the future in birds that are unmuted right after cannula implantation and by inspecting their recovery of singing rate. Presumably, birds were also experiencing an intrinsic burden in the muted state from not being able to sing; however, we are not able to estimate this burden based on our data.

### 4.3. Deafening by bilateral cochlea removal

The zebra finch is a good model system to study the roles of auditory feedback in the production of vocalizations, in learning, and in the neural correlates thereof (Vallentin and Long, 2015).

Deafened birds vocalize for several months after deafening. This response is similar to humans who also continue talking after complete deafening due to disease or accident, provided they have already learned to talk while hearing (post-lingual deafness). Yet, post-lingual deafness in humans leads to speech deterioration (dysphonia and low speech intelligibility) due to long-term absence of auditory feedback (Cowie and Douglas-Cowie, 1992; Lane and Webster, 1991); this side effect of deafening in humans is similar to the deterioration of song found in deafened birds (Konishi, 1964; Lombardino and Nottebohm, 2000; Nordeen and Nordeen, 1992).

Speech deterioration is an important issue for society; post-lingual deaf people losing their job and social interactions often become depressed. Over half a million people in the United States alone suffer from severe to complete hearing loss, leading to high economic costs for the society (Mohr et al., 2000). A common issue in deaf people is problems with vestibular orientation, subject to large inter-individual differences (Braden, 2013; Kaga et al., 2008; Potter and Silverman, 1984). Although most deaf people adapt through increased visual orientation, it is not uncommon for a deaf person to have a severely impaired balance when visual information is limited (e.g: at night).

Deafening birds by bilateral cochlea removal has been used to study the role of auditory feedback in song learning and song maintenance. New insights have been gained, for example by identifying the brain areas responsible for song deterioration after deafening (Horita et al., 2008; Nordeen and Nordeen, 2010; Scott et al., 2007) and determining the structural and functional brain changes resulting from deafening (Tschida and Mooney, 2012).

Deafening is usually done using bilateral cochlea (basilar papilla) removal (Schwartzkopff, 1949; Vallentin and Long, 2015). The surgical procedure for cochlea removal can cause minimal vestibular disturbances. Some studies reported that following cochlea removal, a few birds exhibited opisthotonos-like symptoms (holding the head up and back) along with erratic flight behaviors (birds fell from the perch; they did not reach a perch on first attempt) (Konishi, 1964; Nordeen and Nordeen, 1992). However, these symptoms tended to disappear within 1-2 weeks. (Konishi, 1964) showed that after a short time, deaf birds learned to navigate visually using visual cues in the environment.

For these reasons, it is important not to damage the semicircular canals during cochlea removal and to adhere to strict experiment termination criteria specific to vestibular orientation problems. Following surgery, birds’ welfare should be assessed at 3 different times of the day until their behavior stabilizes. For example, if a bird sometimes misses the perch when trying to land on it, it should at least be able to land on its feet on the cage floor. Such birds with perch-landing problems should be given easy access to food and water (to eliminate problems that could result from low glycemia). If a bird is systematically unable to land on its feet, the experiment should be terminated. It may be possible to define stricter termination criteria using video recordings and automated quantification of behavior.

An alternative method to deafen birds is to chemically destroy hair cells with the ototoxic aminoglycoside. Contrary to mammals, birds can regenerate lost hair cells and hearing recovers after 4 weeks (Woolley et al., 2001). However, because “hair cells in the basal (high frequency) end of the papilla are most vulnerable to chemical ablation, this treatment leads mainly to “high frequency hearing loss”. In a study performed in Bengalese finches, birds could still hear in the 2–6 kHz frequency range following ablation, albeit with a higher auditory threshold. As the fundamental frequency of zebra finch syllables is in the 1–6 kHz range, this method cannot replace cochlea removal to study to role of auditory feedback in vocal plasticity. In addition, there is evidence that zebra finches are not highly susceptible to ototoxic aminoglycoside treatment (DeAngelo, 2008).

One alternative deafening method is to mask auditory feedback with continuous loud white noise. Continuous white noise decrystallizes the songs of adult zebra finches (Leonardo and Konishi, 1999), suggesting that masking noise acts via suppression of auditory feedback, similar to deafening by cochlea removal. Unfortunately, masking noise is also recorded with microphones at the same time as the bird’s song. This co-recording prevents accurate sound measurements, which are required to detect small changes in vocal output for example during song conditioning.

It is difficult to evaluate whether the burden of continuous white noise is smaller or larger than the burden of cochlea removal. Continuous exposure to loud white noise (90–100 dB) in juvenile (40 days old) canaries for 200 days partially destroys hair cells and leads to an increase of hearing threshold by 50–60 dB, and to birds’ inability to perform normal song (Marler et al., 1973). However, when the loud noise was removed, birds improved their song in the following seasons (canaries are seasonal vocal learners)(Marler et al., 1973). Studies on corticosterone release triggered by white noise (60–80 dB) presented during the breeding season did not lead to elevated corticosterone levels in captive zebra finch offsprings, but the noise treatment negatively affected the volumes of some of their song-control brain areas (Potvin et al., 2016). To our knowledge, studies on corticosterone levels and stress indicators in broilers with no white noise experience during youth are missing, but similar studies on poultry showed that noise treatment of 80 dB resulted in a significant stress response (Bedanova et al., 2010).

### 4.4. Backpack for song recording

Animal-mounted sensors have opened the possibility to measure a variety of behaviors in the wild with unprecedented detail. For example, recently developed miniaturized sensors can track animals’ positions throughout several years in real time (Kays et al., 2015).

Until recently, the study of vocal communication was impossible given that audio signals recorded with external microphones do not allow for separation and unequivocal assignment of a vocalization to its producer. For this reason, we and other labs have developed miniaturized sensors suitable for even very small animals (Anisimov et al., 2014; Couchoux et al., 2015; Gill et al., 2016). With these devices, equipped with an accelerometer or microphone, we can now study, for the first time, vocal communication at the individual level.

To be maximally useful, animal-mounted devices should not affect animal behavior. In freely ranging wild birds, studies showed diverse effects on behaviour. In one study, the attached sensor did not lead to negative effects “on the likelihood of breeding, the likelihood of brood desertion, and on nestling-provisioning behaviour” (Snijders et al., 2017). A recent meta-analysis, however, has found that attached sensors in the wild can indeed influence energy investment, behavior, reproduction and/or survival (Barron et al., 2010) in birds. In some cases, the effect of the sensors only becomes evident when the animal was pushed to its physiological limits due to adverse environmental conditions such as cold weather (Snijders et al., 2017; Wilson et al., 2015). Surprisingly, this negative impact seems to be independent of sensor weight (up to 10% of body weight). On the contrary, the location of the backpack on the body is more important than the actual weight. It is advised to position the backpacks close to the animal’s center of gravity (i.e. in the middle to lower back), where their effect on flying is minimized (Vandenabeele et al., 2014).

Contrary to studies in the wild, laboratory birds are held in stable environmental conditions (i.e. constant light-dark cycle, stable temperature and humidity) without predators. Food and water are provided ad libitum and behavior is monitored to take measures to reduce the impact of wearing a backpack (e.g. changing harness size). However, even in captivity, backpacks could impact an animal’s well-being. For example, in the small passerine *Spiza Americana*, within 24 h after transmitter attachment in captivity, corticosterone levels increased, but returned to baseline levels within two days (Wells et al., 2003). Therefore, as corticosterone returns to baseline, aversive effects of backpacks are likely caused by the adaptation to wearing the backpacks.

In our lab, zebra finches seem to experience some initial discomfort upon fastening the backpacks (e.g. birds pick on the backpack). In addition, for heavy backpacks (~20% of body weight), locomotor activity, as well as singing and calling rates, are transiently reduced, but return to baseline within 10 days after mounting of the backpack (Anisimov et al., 2014). With lighter backpacks (<10% of body weight), the accommodation occurs even faster, within 3 days (Gill et al., 2016). Given that the average zebra finch in our colony weighs ~16 g, our backpacks (1 to 1.5 g) correspond to between 6.3% and 9.4% of body weight. Although our backpacks are heavier than devices recommended for usage in the wild (<5% of body weight), small birds are able to carry a higher load than heavy birds (Caccamise and Hedin, 1985). Birds with backpacks perform all normal behaviors including grooming, calling/singing, flying, hopping, eating, drinking, and copulation (Anisimov et al., 2014; Gill et al., 2016). Judged by visual inspection, it seems that birds stop noticing the backpacks after a while, similar to humans that stop thinking about the clothes they wear.

To conclude, the establishment of backpacks is opening new possibilities to study vocal communication with individual-level resolution in a social setting (Anisimov et al., 2014; Gill et al., 2015; Stowell et al., 2016). Currently, the initial backpack attachment and battery exchange exert a transient reduction on locomotor activity and vocalization. Although we might not be able to fully eliminate the effect of handling, future advancement of battery technology and electronics will allow us to keep reducing the backpack’s weight and, in this way, the burden onto the birds.

## 5. Recordings of neural activity

In-vivo electrophysiology has been the most successful technique for measuring the role of single nerve cells in generating motor behavior. Intracranial fine electrodes provide a readout of neuronal activity with very high spatial and temporal resolution, unlike readouts provided by non-invasive techniques such as functional magnetic resonance imaging (fMRI), electroencephalography (EEG), or magnetoencephalography (MEG). In-vivo electrophysiology has been the main driver of recent neuroprosthetic advances, allowing paralyzed patients to regain arm reaching and hand grasping functions (Chapin et al., 1999; Nicolelis, 2003; Taylor et al., 2002; Wessberg et al., 2000), and vocal production (Guenther et al., 2009) using brain-machine interfaces. Single neuron and population level recordings are important tools for neurosurgeons trying to locate the focal points of epileptic seizures in patients (Blume et al., 1997). To reveal the underlying mechanisms that generate epileptic states in the brain, scientists resort to animal models (Grone and Baraban, 2015).

Our work of measuring neural activity in singing zebra finches using fine metal electrodes mounted on miniature microdrives has led to the discovery of a representation of time in the brain (Hahnloser et al., 2002). When this song-control brain area is cooled, the song slows down (Long and Fee, 2008). When the analogous brain area in humans is cooled, the duration of spoken words in humans changes (Long et al., 2016). As a consequence, this line of songbird research has brought benefits to epileptic patients by simplifying procedures used to localize speech areas during therapeutic surgery (Flinker and Knight, 2016).

### 5.1. Recordings during unrestrained behavior

All natural behaviors, such as exploration of a space through locomotion (O’Keefe and Dostrovsky, 1971), vocalization (McCasland, 1987), and social interactions (Katayama et al., 2013) are controlled by electrical activity in populations of neurons. Experiments in unrestrained, freely-moving animals have been key for understanding these neural control mechanisms. Typically, in these experiments, small electrodes are chronically implanted in the brain, permitting the recording from individual neurons or clusters of neurons while animals move around with relatively little constraints. For example, a series of seminal electrophysiological studies have shown that the brain creates a map of the external space. Namely, the brain controls navigation through a complex environment by means of cells that react to a specific place (“place cell”) (O’Keefe, 1976) or to a structured lattice of places, encoded in the activity of so-called “grid cells” (Fyhn et al., 2004). These studies have lead to the Nobel Prize in Physiology and Medicine in 2014.

In typical electrophysiology experiments, metal electrodes, carbon fibers, or glass pipettes are implanted into the brain in close proximity to the targeted cell types. In birds, the electrodes can be mounted on a chronically implanted electrode holder or microdrive (Fee and Leonardo, 2001). When microdrives are equipped with a motor, the electrode tip can be repositioned in search of new neural signals without having to handle the animal, which can be a stressor that suppresses the behavior of interest (very often, neural signals are lost during a recording because the brain tissue moves relative to the electrode, in many cases neurons cannot be recorded for more than 10 minutes). Current state-of-art devices using feedback control mechanisms allow a maximum movement range of 6 mm within which electrodes can be positioned with a guaranteed precision better than 1 μm (Jovalekic et al., 2017). This range allows for targeted recordings from single neurons within an extended network of brain areas.

Songs produced before and after implantation surgery are visually indistinguishable (Fig. 5-1). One obvious burden of electrophysiology for animals is that the implants and its attachments (e.g. microdrives, headstages, and connectors) add weight to the animal’s head. As discussed below section, 10% of the animal’s weight can be an upper limit for the total weight of an implant. Mechanical/electrical microdrive designs have improved over the years and motorized microdrives have become much lighter, weighing less than 1 g (Jovalekic et al., 2017), which amounts to less than 10% of even the lightest adult zebra finch. At the same time, to better decode the behavioral intentions of a subject and to discount for intrinsic noise in single-neuron activity, it is becoming clear that large populations of neurons must be recorded (Harris et al., 2016). This means that electrophysiologists must introduce higher numbers of electrodes into the brain. To not impact animal welfare, the additional weight resulting from the additional electrodes must be compensated by improved microdrive designs. Thus, important advances in animal welfare can still be achieved through ingenious mechanical and electrical engineering. One strategy to improve welfare could be to distribute the weight among the head and the back, as demonstrated by telemetric recording systems in which the batteries and/or the telemetry circuits are placed on the back (Hasegawa et al., 2015; Liberti et al., 2017).

**Figure 5-1.**
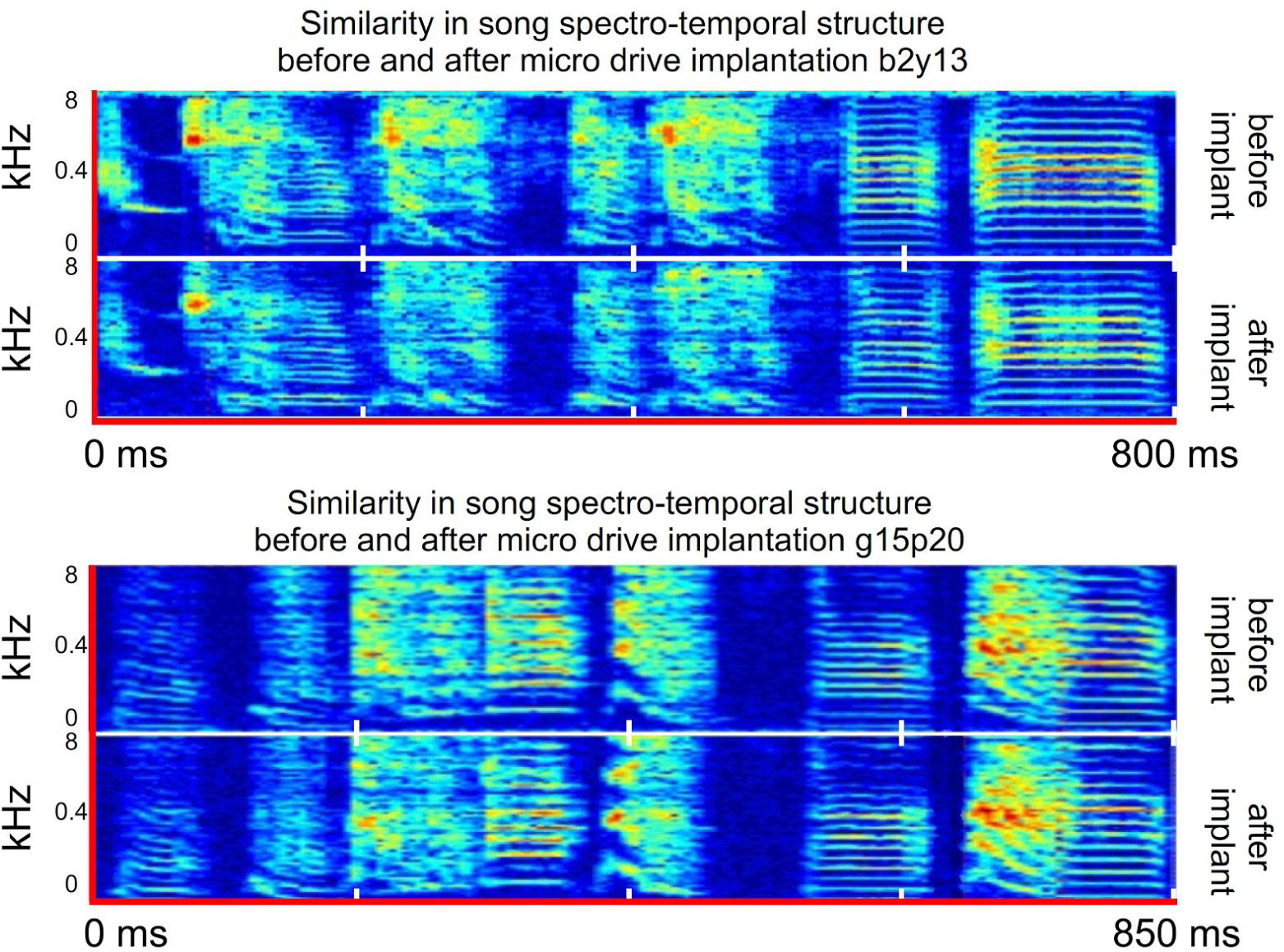
Songs before and after implantation surgery are highly similar. In two animals, shown are logarithmic sound spectrograms of two song motifs, one produced before surgery (top), and the other after surgery (bottom). They are visually indistinguishable.

#### Tethering

To deliver the electrical signals from the microdrive to the recording system, the implanted device is connected to a tether cable. The tether cable can limit birds’ locomotion by adding tension and/or torque on the head, potentially leading to unnatural head postures. Some of these problems can be counteracted by use of a motorized commutator that cancels out the tether’s torque along the gravitational axis. The commutator allows minimally constrained head movements thanks to a magnetic sensor and electrical motor that continuously rotate the commutator into a torque-free position (Fee and Leonardo, 2001). Because of tethers, perches should be placed at very low positions to avoid animals trying to pass underneath and getting entangled. In spite of use of commutators, tethers limit cage size as long tethers can lead to entanglement of animals. Tethering currently seems to be prohibitive for studies of free-ranging behaviors such as foraging, breeding, pair formation, and vocal communication across large distances, just to mention a few.

Tethers can be replaced with a wireless telemetry system (Schregardus et al., 2006). Two standard hearing aid batteries (zinc–air type) can power a telemetry recording device for 15–22 h, but current telemetric devices provide only a single electrode for neural recording and no possibility for electrode repositioning. Given current technology for multichannel recordings, a 1 g battery (e.g. lithium-polymer rechargeable battery) cannot provide enough energy for several hours of recording. Battery replacement requires frequent handling, disrupting the birds’ natural behavior and suppressing its singing rate (Gill et al., 2016). Thus, tethering is mandatory if long-term recordings or recordings with multiple electrodes are required and animal handling to be avoided.

#### Burden of weight load on the head

The implants for brain activity manipulation and recording (microdrives, headstages, head plates, connectors, etc.) add weight to the animal’s head. Schregardus and colleagues (Schregardus et al., 2006) studied the short-term effects of anesthesia, surgery and implantation of a 1 g telemetric neural implant. They show that the amount of hopping and singing duration is severely reduced post-surgery in zebra finches that are anesthetized compared to those that are simply handled. However, there was no statistically significant difference in hopping and singing rates between birds that were anesthetized and implanted with the device and those that were simply anesthetized without surgery. Additionally, compared to baseline, hopping and singing rates were only reduced dramatically on the day of and day after surgery, recovering to baseline rates within 2 days.

To our knowledge there is no study quantifying the long-term effects of carrying an ~1 g weight on the head of small animals. In our experience, zebra finches usually stand erect and keep their usual posture for the implant weighing up to 10% of the animal’s weight (up to 1.6 g for an average zebra finch in our colony weighing ~16 g) as discussed in the use of backpack (see Section 4.4). We propose to use the beak tracking system (as described in Section 2.1) to check how birds adapt to the 10% weight after implantation and recover their posture, singing rate, and locomotion activity. A weight reduction device that pulls up the bird’s head by a thin string may be useful to support the bird’s posture and locomotion, similar to one used in rodents (du Hoffmann et al., 2011).

Unfortunately, our data currently does not allow us to estimate the cumulative burden of tethering and of wearing an implant, because we have not been recording songs around the clock in these experiments. To save hard disk space, we saved songs to disk only when a good neural signal was present on the electrode. In future studies, it will be desirable to monitor singing rate and movement behavior before and after tethering, in order to obtain more information about the impact of chronic neural recordings on zebra finch welfare.

### 5.2. Head fixation for optical imaging

What is the burden of daily head fixation for high-resolution optical imaging?

In the last decades, our understanding of brain and behavior has progressed thanks to the development of new recording techniques. One of these, two-photon laser scanning microscopy, allows for longitudinal recording of neural activity from population of neurons over periods of several months (Crowe and Ellis-Davies, 2014; Li et al., 2017; Sadakane et al., 2015). Two-photon imaging, most frequently used in mice, has recently been applied also to zebra finches to reveal drastic structural changes in synapses triggered by exposure to tutor song in circuits responsible for song motor control (Roberts et al., 2010), and to study the correlational structure of auditory responses to song stimuli (Graber et al., 2013; Peh et al., 2015). Also, thanks to the large number of neurons that can be simultaneously recorded, two-photon imaging enabled the discovery of a critical population of song premotor neurons that encode song time in a continuous rather than discrete manner (Lynch et al., 2016; Picardo et al., 2016). This latter discovery is of particular relevance because the neurons of interest are the only premotor types that undergo neurogenesis in the adult bird, a property that is believed to hold enormous therapeutic potential if exploitable in humans (Goldman, 2016; Lindvall et al., 2004)

Two-photon microscopy is less invasive than microelectrode recordings, because fluorescence signals can be imaged through a transparent cranial window, and in principle no brain penetrations are required for imaging, only a craniotomy. Imaging is subject to less spatial restriction than electrophysiology, and is more suitable for longitudinal studies of motor learning and neural network dynamics (Crowe and Ellis-Davies, 2014).

Two-photon microscopes are very large and heavy, which is why imaging must be performed in the head-fixed state. Head-fixed birds can be induced to sing (78/109 animals tested) when using a training procedure with reward delivery (Picardo et al., 2016). Miniaturized two-photon microscopes for freely moving studies have been developed (Flusberg et al., 2005; Helmchen et al., 2013) and successfully applied in rats and mice (Helmchen et al., 2013; Sawinski et al., 2009). However, the excessive weight of such devices (>3 g) is currently prohibitive for use in lightweight animals such as zebra finches. Therefore, there is currently no alternative method for high-resolution imaging of neural activity other than head-fixation under a two-photon microscope.

For rapid and flexible head fixation, lightweight head plates have been developed that are chronically implanted and allow birds to be head fixed with short handling times. Typically, titanium head plates used in zebra finches weigh less than 0.5 g. In order to reduce excessive escape reactions and injuries while head fixed, birds are kept in a size-adjustable restraining jacket (using velcro) for up to 2 hours per imaging session. Scapulars are restrained and the wings protrude from holes in the jacket. No signs of injury or abrasion on the body have ever been observed following the use of such jackets. Restraining the body for two hours is expected to lead to increased corticosterone levels (Banerjee and Adkins-Regan, 2011). Restrained animals tend to rarely vocalize. Recently, it has been shown that head-fixed but unrestrained zebra finches standing on a perch can have a natural body posture and produce normal songs (Picardo et al., 2016). It seems that removal of the restraining jacket and preparation of animals to head-fixation using training are refinements that could be worth adopting in future studies.

Social interactions increase vocal learning success (Chen et al., 2016). To allow for visual and vocal interactions with a companion bird during imaging, we provide ambient light near the birds. During the 2 h imaging session, a female bird and/or tutor is introduced at a distance of less than 20 cm from the head fixed animal. Live tutors typically sing 10 to 30 songs during a 2 h session, which seems to be the minimal condition to achieve successful tutoring and statistically meaningful data analysis (Tchernichovski et al., 1999). Additional introduction of a female in a separate cage facing the tutor increases the number of tutor songs to 20–50 during the 2 h imaging session.

Care must be taken that animals in such head-fixed imaging experiments get enough sleep, food, and rest in between imaging sessions. In a preliminary study we monitored animals’ vocalization rates and body weight in resting periods following each imaging session. Because young birds do not produce recognizable song motifs, we simply counted the number of vocalizations including song syllables and calls.

We found that animals gradually increased their vocal output after head plate implantation (Fig. 5-2A, B). Initial head fixation and imaging did not markedly decrease vocal output during resting periods, suggesting that birds coped well with head fixation. Over the course of the experiment, animals almost tripled their vocal output (Fig. 5-2A). The body weight of head-fixed juveniles was within the range of normal body weight of 18 age-matched control birds from our colony (Fig. 5-2C). In one bird there was a small tendency of weight “rebound” after release from daily imaging (Fig. 5-2C), but it is not clear to what extent this weight increase can be attributed to either release from head-fixation or the continual growth.

**Figure 5-2.**
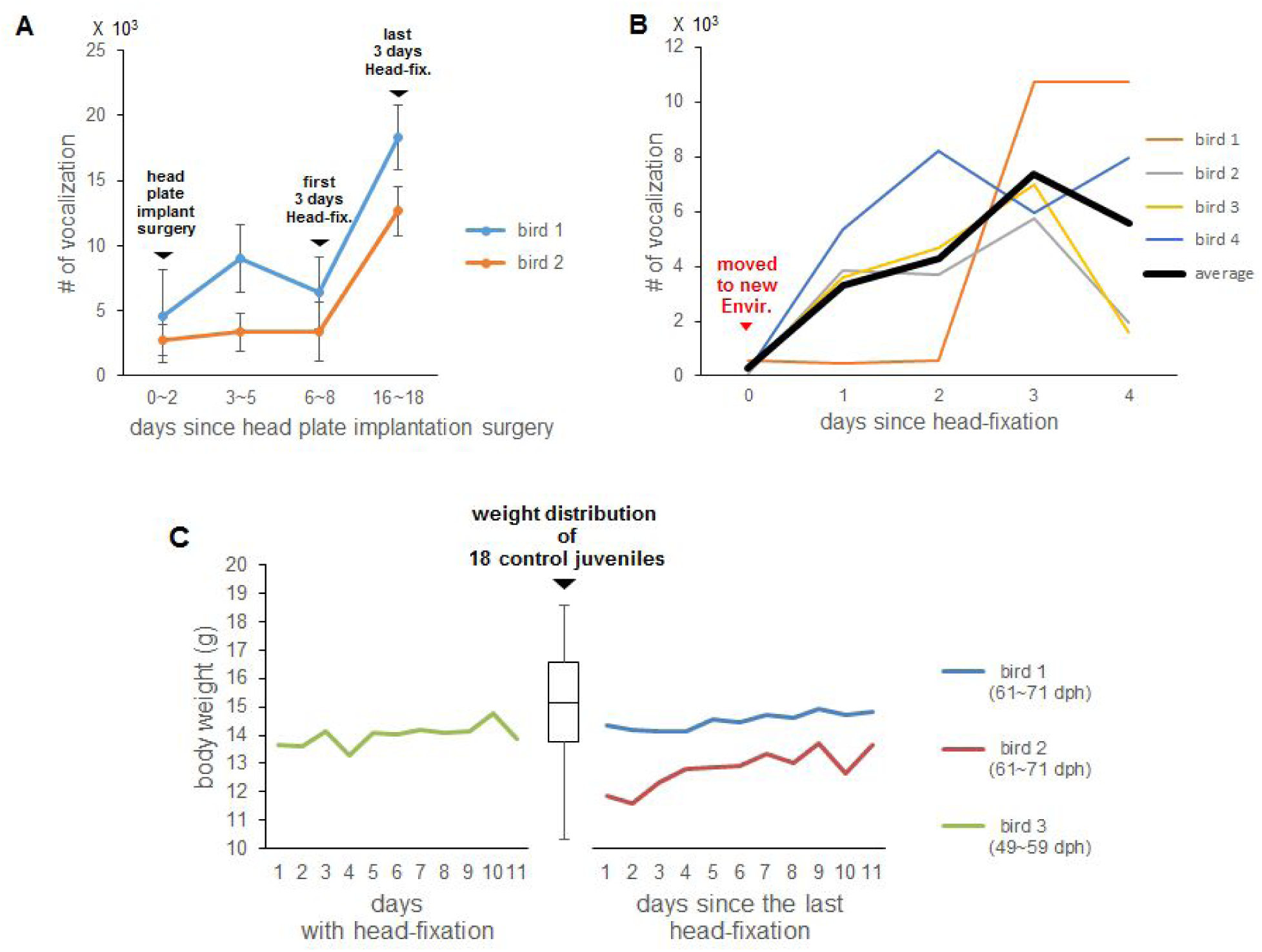
Behavioral monitoring in head-fixed juveniles. (A) Number of vocalizations during resting periods. After day six, birds were head fixed 1 h in the morning and 1 h in the evening each day. Each point represents the number of vocalizations averaged over 3 days in the same condition. Error bars indicate standard deviations across the 3 measurement days. (B) Number of daily vocalizations produced on 5 days with 2h of head-fixation per day (data from 4 animals shown in color, the average vocalization rate is shown in black). Animals were subjected to the cumulative burden of head plate implantation on day -1, 2) environment change starting on day 0, and 3) 2h daily head fixation between days 0 and 4. Vocalization rates gradually recover. (C) Body weight of juveniles during (left) and after (right) head-fixed imaging period. Box-and-whisker diagram represents weight distribution of 18 juveniles from our colony (reared by father and mother, not used in experiment, age range 43–76 dph): Min. (bottom whisker), first quartile (bottom of the box), median (center line of box), third quartile (top of the box), and max. (top whisker) body weight. Bird 1 and Bird 2 were head-fixed for 11 days (2 h/day) before the body weight measurement and not head-fixed during the 11 days of body weight measurement. Bird 3 was head-fixed daily for 2 hours during the 11 days of body weight measuring period. All three birds were separated from the father at 15 dph, reared by their mother during 15–35 dph, and isolated individually in soundproof boxes from 35 dph until the end of the experiment (115 dph).

Songs produced in the head-fixed state are indistinguishable from songs produced during free movement (Picardo et al., 2016). Thus, there is no reason to believe that the neural mechanisms of vocal production studied in manipulated birds (subjected to surgeries for virus injection, head-post implantation and head-fixation) are to be different from the mechanisms at work in the healthy and unmanipulated brain.

The cumulative burden in these experiments (isolation in a new environment, surgery, and head-fixation), as estimated using body weight measurements, seems to be non negligible. Unfortunately, we currently do not have any data for evaluating the burden based on vocal output. The number of vocalizations remain stable in the first 2 days after head-plate implantation, which may indicate either that the surgery had little impact on the bird’s welfare or that it only recovers very slowly from the adverse effects of the surgery. Birds seem to cope well with daily 2 h of head-fixation, evidenced by the strongly increasing number of vocalizations they produce in resting periods following the imaging sessions (Fig. 5-2A).

Based on the work of (Picardo et al., 2016), improvements of our procedures to minimize stress and optimally prepare animals to head fixation could be to train non-head fixed animals to remain on a particular perch or in a small quadrant of their cage to obtain food or social rewards.

## 6. Conclusion

In the present study, we have discussed what is known about stress caused by procedures used in research on zebra finches. Wherever possible, we have provided our own data and proposed possible refinements.

Based on our findings we suggest that singing rate might be a useful readout of welfare. Because singing is the object of interest in many laboratories, this welfare readout could be adopted with little effort in adult males. To quantify song output in juveniles is harder and will require efforts, mainly focused on the question whether to include or exclude some calls in the analysis and whether these calls can be automatically detected in juveniles.

Our investigation of isolated birds prompted us to draw the intriguing conclusion that isolation is not stressful to zebra finches, but isolation in a new environment is. Our data to this effect could not have been more revelatory: Birds isolated in a new environment immediately suppressed their singing for some time, whereas birds isolated in their current environment immediately increased their singing (Fig. 2-6). Because the increase in song output was immediate and stable in the latter animals, we infer that birds were highly motivated to sing and that the increase was not due to a gradual maladaptation to an environmental stressor. Although it is not clear whether there can be a point at which too much singing should be considered pathological, there is currently no data known to us that would suggest so.

Besides singing rate, our proposed automatic beak tracking system could be routinely implemented as an independent method to monitor movement behavior in a variety of conditions (Section 2.1). Further improvements of this technology might lead to automatic detection of abnormal behaviors such as stereotypies. Of course, many other parameters should be considered for assessing the welfare of zebra finches in experiments. However, there is currently a lack of such data from our lab.

In summary, although all the discussed procedures induce some level of acute burden, birds usually adapt to the considered manipulations within a few days (i.e. they recover singing rate and show minimal weight loss). Therefore, adaptive coping mechanisms in zebra finches might be in place to rapidly reduce stress levels.

## Acknowledgement

We would like to thank Annamari Alitalo for helpful comments about the manuscript.

## Materials and Methods

### Animals

Data from a total of 132 zebra finches (*Taeniopygia guttata*) were used in the present article. Birds were obtained from our breeding colony or from a local supplier. In the colony, they were in constant social contact. All birds were maintained on a 14/10 h day/night schedule and were given ad libitum access to food and water. All procedures described here were approved by the Veterinary Office of the Canton of Zurich, Switzerland.

### Song recording and syllable labelling

For song recording, birds were housed in soundproof boxes. Songs were recorded with a wall-attached microphone (Audio-Technica PRO42) at a sampling rate of 32 kHz into computer hard disks, and analyzed with a custom script (*FlatClust*) implemented in the MATLAB software (Mathworks Inc). To visualize songs, we computed log-power sound spectrograms, which are time-frequency representations of sound intensity. Song syllables, calls, and noise (e.g. wing flapping) were first separately labelled by visual inspection of the spectrograms for a small subset, and then, automatically discriminated by a perceptron algorithm. The number of song motifs was calculated by counting a specific syllable (or a part of the syllable) always present in a song motif. In case of multiple birds housed in the same soundproof box, labelling and counting were manually corrected to untangle overlaps of vocalizations with visual inspection of spectrograms.

### Song similarity analysis

Male juvenile zebra finches (< 90 days; *N* = 12) used to study the effect of housing condition on song development (in Section 3.1) were raised in our local animal facility. Juvenile birds were isolated from father from 15 days post-hatching (dph) on, and were fed by the mother alone until 35 dph. The male juvenile birds were then transferred to individual cages in soundproof boxes. Then, we divided the juvenile birds into three groups and tutored them until 59 dph as follows. Group 1: the juvenile bird was housed together with an adult male (24 hours per day); Group 2: the juvenile was housed together with an adult female placed in the same cage as the juvenile bird outside the tutoring period. For 90 minutes every morning, the female was exchanged with an adult male; Group 3: the juveniles was housed alone in the soundproof box and an adult male was presented for 90 minutes per day starting on 46 dph. Each group included four birds.

For each bird, 20 pupil (juvenile) song motifs and 20 tutor song motifs were manually selected from the last day of recording (59 dph of the juvenile bird). Pair-wise comparison of the pupil songs and tutor songs were performed with Matlab and Sound Analysis Pro software (SAP 2011) (Tchernichovski et al., 2001) in batch comparison mode. The resulting 400 comparisons were averaged and a mean similarity score was generated for each tutored juvenile bird.

We analyzed statistical significance in similarity differences among the three tutoring methods by a linear mixed model. We used the model to evaluate whether the fixed effects that characterize differences among tutoring groups significantly deviate from zero. For birds in the 24 h/day tutored group (Group 1), we modeled the song similarity *y*^24*h*^ as the sum of a fixed (non-zero) mean group similarity β^24*h*^, a random individual bird effect *b*^24*h*^, and a random observation error: ε ^24*h*^: *y*^24*h*^ = β^24*h*^ + *b*^24*h*^ + ε^24*h*^. For the birds in one of the 90 m/day tutored groups (Group 2), we included in the linear model a fixed bias term β^90*m*^ relative to the average similarity β^24*h*^ of Group 1 : *y*^90*m*^ =β^24*h*^ + β^90*m*^ + *b*^90*m*^ + ε^90*m*^. Our statistical tests then consisted of evaluating whether β^90*m*^ > 0 for either of the 90 min/day groups (with and without female).

### Body weight measurements

The body weight data were assembled from health inspection procedures in various experiments. Each bird was weighed between 8:00 to 11:00 to avoid variations due to circadian rhythms (Dall and Witter, 1998). The weighting procedure (Boyd, 2012) takes in total 0.5 to 2 min.

### Beak tracking

We have developed a custom-made software for real-time tracking of the color of the bird’s beak. Adult zebra finches have a characteristic red (male) and orange (female) beak color, thus their beaks can be used as an identifiable marker to detect each bird’s location and movement. In a preliminary test, one tethered bird was monitored by our tracking system at the same time of neural recording (the tracking did not add extra burden onto the bird). A USB webcam (C905, Logitech) was position on the top of cage, and top-view images were obtained at 30 frames/s with a spatial resolution of 640 × 480 pixels. The software detected pixels that have the salient beak color by masking with a predefined HSV color key (hue: 0–30, saturation: 50–100, value: 20–100). Then, the median location of detected pixels was calculated in each image. If the number of detected pixels was less than 40, the location was not calculated and set to zero. The calculated locations were saved into a data file with timestamps. Two types of tracking maps were generated. The position map shows the spatial density of the animal, and the motion map (a summary of movement paths) shows the most frequent paths taken by the animal. Firstly, the missing data points were interpolated by just filling previous data points since we assumed the bird was located at the same place. Then, the position map was generated as a two-dimensional histogram with bins of 8 pixels. For generating the motion map, we calculated lines between two location points in two successive images and accumulated the lines into a two-dimensional histogram with bins of 8 pixels. Lines in which the lengths were less than 30 pixels were excluded from the accumulation in order to remove tiny fluctuations in the tracking data. To evaluate similarity between maps obtained from two different time period we used Pearson’s correlation coefficient.

### Statistics

All statistical tests were performed with a significance level at *α* = 0.05. If we compared several data points for each bird before and after a manipulation, we estimated the change in percent by dividing the data points after manipulation by the mean from before the manipulation. We tested whether the data points after the manipulation (in percent) significantly differ from one using a t-test to show whether there is a significant change from before to after manipulation.

